# Multi-scale dynamical modelling of T-cell development from an early thymic progenitor state to lineage commitment

**DOI:** 10.1101/667709

**Authors:** Victor Olariu, Mary A. Yui, Pawel Krupinski, Wen Zhou, Julia Deichmann, Ellen V. Rothenberg, Carsten Peterson

**Affiliations:** Computational Biology and Biological Physics, Lund University, Lund, Sweden; Division of Biology and Biological Engineering, 156-29, California Institute of Technology, Pasadena, California 91125, USA

## Abstract

Thymic development of committed pro-T-cells from multipotent hematopoietic precursors offers a unique opportunity to dissect the molecular circuitry establishing cell identity in response to environmental signals. This transition encompasses programmed shutoff of stem/progenitor genes, upregulation of T-cell specification genes, extensive proliferation, and commitment after a delay. We have incorporated these factors, as well as new single cell gene expression and developmental kinetics data, into a three-level dynamic model of commitment based upon regulation of the commitment gene *Bcl11b*. The first level is a core gene regulatory network architecture determined by transcription factor perturbation data, the second a stochastically controlled epigenetic gate, and the third a proliferation model validated by growth and commitment kinetics measured at single-cell levels. Using expression values consistent with single molecule RNA-FISH measurements of key transcription factors, this single-cell model exhibits state switching consistent with measured population and clonal proliferation and commitment times. The resulting multi-scale model provides a powerful mechanistic framework for dissecting commitment dynamics.

## INTRODUCTION

Hematopoietic progenitors continually replenish the body’s supply of blood cells including T lymphocytes. Many of the important regulatory factors controlling T-lineage commitment are known (reviewed in^1,2^), as well as their patterns of expression^3-5^. Specific targets of several of these factors have been defined by perturbation studies, providing a strong basis for understanding the dynamic operation of an intrinsic gene regulatory network (GRN)^6-26^. However, the timing of observed transitions and evidence for a rate-limiting role of epigenetic constraints prevent modeling of the commitment process through a simple factor-target interaction approach. This paper presents a three-level model that integrates current understanding of this intrinsic GRN with insights into epigenetic control points and population dynamics characterizing T-cell commitment.

Small numbers of multipotent hematopoietic precursors migrate continuously from bone marrow into the thymus^27,28^ where they enter the T-cell developmental pathway, driven by Notch signalling and cytokines in the thymic microenvironment. These early progenitor cells are CD4 and CD8 double negative (DN), as well T-cell receptor negative. The Kit^high^ early thymic progenitors (ETP or DN1) proliferate before transitioning to DN2, a step marked by CD25 surface expression, then upregulate *Bcl11b* and undergo Bcl11b-dependent T-lineage commitment (Fig. 1A)^13,14,24,29,30^. Expression of the zinc finger transcription factor *Bcl11b* not only alters many aspects of genomic activity and cell function^24,31,32^, but also correlates with the functional committed state of single DN2 cells^24^ and can thus serve as a proxy for commitment. After *Bcl11b* activation Kit^high^ DN2a cells move through DN2b into the Kit^low^ DN3 stage, after which they can continue differentiation after rearranging a signalling competent T-cell receptor (Fig. 1A). A broad shift in expression of multiple regulatory genes occurs during commitment, and the stem/progenitor transcription factors expressed in DN1 and DN2 stages are generally extinguished (reviewed in^1,33^).

**FIG. 1.**
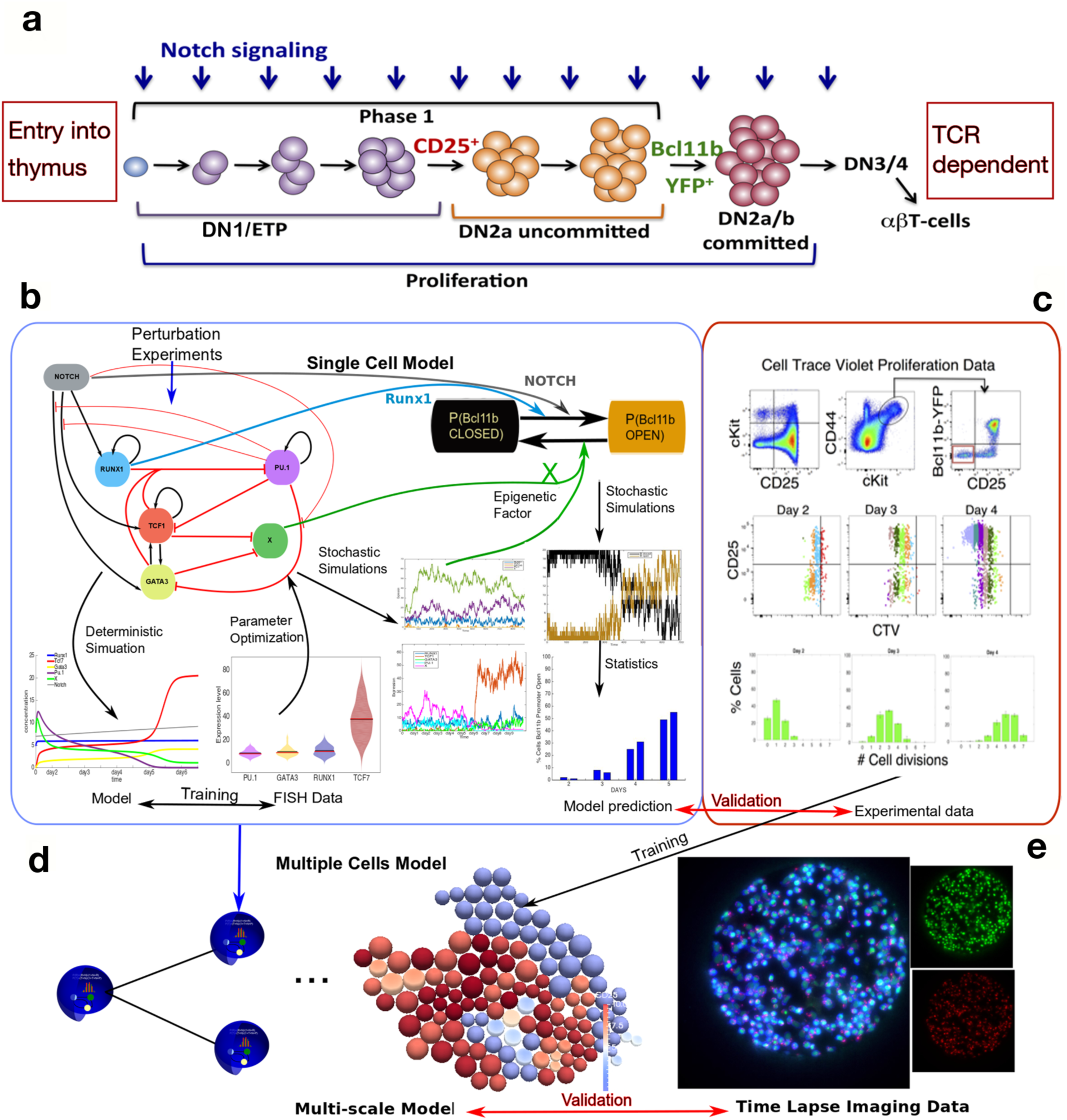
Multi-scale modelling roadmap. **A** Schematic of T-cell development. Figure shows initial Notch-dependent stages after multipotent precursors immigrate to thymus. The most immature Kit^high^ Early T Precursors (ETP) are referred to here as DN1. At DN3, T-cell receptor gene rearrangement occurs, and all subsequent development depends on T-cell receptor expression and signalling. **B** Bipartite model for control of onset of *Bcl11b* expression in single cells. **B-left** deterministic component of model: establishing permissiveness for *Bcl11b* activation. **B-right** stochastic component of model: probabilistic onset of *Bcl11b* transcription via positive transcriptional regulators and removal of local epigenetic constraint. Figure shows flow chart of research in this paper. The deterministic model parameters are optimized by fitting simulation results to single cell FISH data. X activity levels represent a constraint function for activity of the *Bcl11b* regulatory region in the second-level, epigenetic stochastic computational model (green arrows). Stochastic simulations of the resulting single cell multi-level model are conducted and statistics of percentage of cells opening the *Bcl11b* regulatory region at each day of experiment are predicted. **C** Experimental measurements of proliferation and developmental kinetics from DN1 to committed T-cells. The number of cell divisions is measured using Cell Trace Violet, as well as the percentage of CD25-positive and Bcl11b-YFP-positive cells, for validation of the single cell multiple stochastic simulation predictions (red arrow). **D** The single cell model is subsequently plugged into the multi-cell population dynamic model (blue arrow) using parameters trained with empirical proliferation data. Predictions from the multi-scale model are finally validated (red arrow) using **E** live-cell time-lapse imaging experimental data measuring proliferation and differentiation of individual DN1 clones.

An important question is how the T-cell commitment decision is controlled. Computational modelling of GRNs represents an important approach for studying mechanisms controlling cell-fate decisions^34^. Here, we have developed a GRN model that incorporates the developmental period from thymic entry by multipotent progenitors to *Bcl11b* up-regulation and T-lineage commitment. Further, new experimental measurements have defined the relationship between proliferation timing and differentiative progression. These have enabled us to integrate this GRN model into a population-level model, as a framework for revealing new factors and roles for key known lineage commitment factors and to conduct dynamics analyses at both single cell and population levels for identifying transient states and general rules for cell differentiation.

Earlier dynamical models dealt with the flux of T-cell progenitor populations through the DN1 to DN4 stages^35^ and the transcriptional network for *Bcl11b* activation assuming deterministic, direct transcriptional regulation by positive regulators Notch, GATA3, and TCF-1^36^. Both models incorporated experimental evidence, but when juxtaposed, a contradiction emerged. The population model^35^ showed that primitive T-cell precursors transit from DN1 to DN2 after 7-10 cell divisions *in vivo*. Activation of *Bcl11b* takes place even later, 2-4 days after the cells become CD25^+^ DN2s *in vitro*^24^. However, all components of the transcriptional GRN model^36^ are expressed in the DN1 stage. A simple combination of previous models would predict *Bcl11b* to be turned on multiple cell cycles earlier than it is, well beyond the range of measurement error, suggesting that the deterministic assumption about *Bcl11b* control itself was incorrect or incomplete.

Incorporating a separate, stochastic timing step downstream of a deterministic network could reconcile these features. In fact, recent evidence implies that the timing discrepancy reflects an intervening epigenetic event. Extensive repression marks and methylation are removed from the *Bcl11b* locus during its activation^37,38^, and the two alleles of *Bcl11b* within the same DN2 cell nucleus can become activated at different times, with discordances of up to 2+ days (and several cell cycles) despite exposure to the same transcription factors^39^. Consequently, in the model proposed here, an additional stochastic timing event for epigenetic activation has been interposed downstream of the action of the initial trans-acting regulators. This additional stochastic step, governed by a new composite parameter, X, and opposed by positive inputs Runx1 and Notch, permits the model to mimic the observed biological time course while using realistic rate constants and parameter values for the regulatory factors that agree with measured values in single cells.

Using these published data and our new experimental results, we propose a three-stage dynamic model of early T-cell commitment. The gene network aspects of the model take into account the unique linkage of commitment to *Bcl11b* activation, and encompasses transcription and epigenetic levels of *Bcl11b* regulation to account for its proximal activation kinetics (Fig. 1). Step 1 is a single cell GRN model based on previously published perturbation experiments. Step 2 is an epigenetic model to account for local chromatin constraints limiting *Bcl11b* activation, even after trans-factor requirements are met. The GRN model parameters were trained on new gene expression measurements from single molecule RNA-fluorescent in situ hybridization (smFISH). We have then embedded the two-step single-cell gene network model into a population model, trained with newly precise cell culture dynamics data, and validated by time-lapse imaging data of cultured DN1 clones. The resulting multi-scale model reveals mechanisms controlling early T-cell development and enables dissection of the kinetic controllers of commitment.

## RESULTS

### Experimental Results

#### Proliferation and Development from DN1 (ETP) to Commitment

Cells already within the DN2 stage, but still Bcl11b-nonexpressing, require ∼2 days on average for *Bcl11b* activation^24^. However, to relate the output of committed cells to events potentially occurring days earlier, we required new benchmark measurements of proliferation and development kinetics from DN1 to T-lineage commitment, and in particular the intervals between the DN1-DN2 transition and *Bcl11b* activation in individual clonal lineages. Bulk and clonal cultures were therefore analysed, starting with DN1 cells purified from thymi of Bcl11b-YFP reporter mice^24^ (Fig. 2A). Sorted DN1s were stained with Cell Trace Violet (CTV) dye that tracks each cell’s proliferation history through dilution, and then co-cultured with Notch-ligand-expressing OP9-DL1 stromal cells and growth-supporting cytokines, IL-7 and Flt3L, to promote T-lineage development. Parallel DN1 cultures were harvested after 2—5 days and assessed by flow cytometry for total cell numbers, CTV-fluorescence intensity, the developmentally-regulated surface marker, CD25 (marking transition to DN2), and Bcl11b-YFP (Fig. 2B). The number of divisions each cell had experienced was determined by flow cytometric measurement of residual CTV staining as calibrated in control cultures (Fig. S1). Numbers of cell divisions experienced by individual cells are represented as different colours in flow cytometry plots (Fig. 2B) and as population distributions of cells with different numbers of cell divisions at each time (Fig. 2C).

**FIG. 2.**
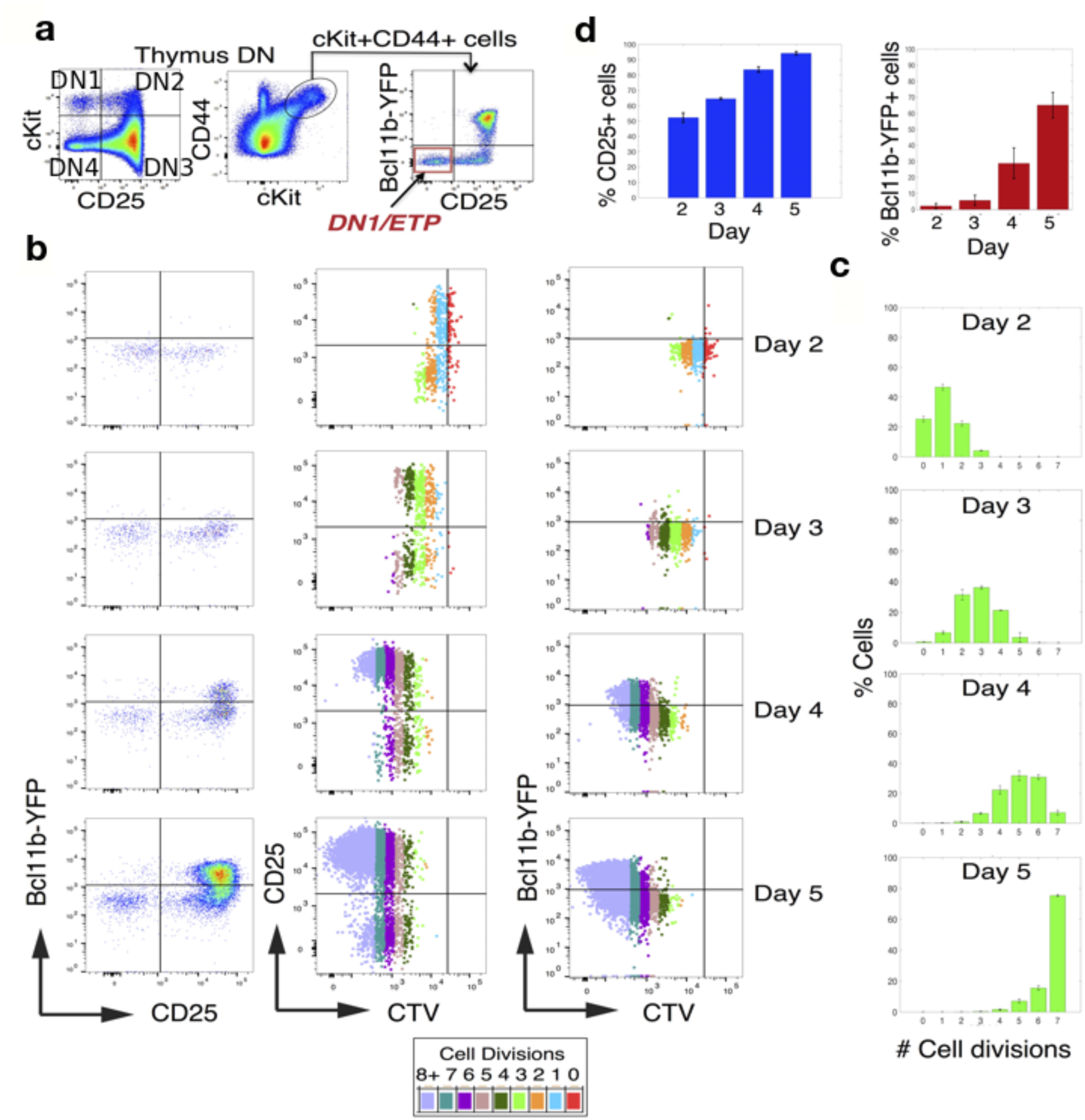
Proliferation and developmental kinetics from multipotent DN1 progenitors to T-lineage committed cells. **A** Flow cytometry plots showing early T-cell developmental stages (DN1-DN4) and the gating strategies used for purifying progenitor DN1 cells, Kit^high^ and lacking CD25 and Bcl11b-YFP, for *in vitro* experiments. **B-D** In vitro analysis of proliferation and development from sorted thymic DN1 cells. DN1 cells were purified from thymi of Bcl11b-YFP reporter mice, stained with a proliferation tracking dye, Cell Trace Violet (CTV), co-cultured with Notch ligand expressing OP9-DL1 stromal cells, and harvested for flow cytometric analysis on days 2-5. **B** Flow plots show the upregulation of CD25 followed by Bcl11b-YFP expression from days 2 to 5 (left), and the relationships between CD25 (middle) and Bcl11b-YFP (right) expression and the numbers of cell cycles each cell has experienced as determined by CTV levels (colour coded by cell division as shown; see Fig. S1 for details). **C** Summary plots of the distributions of the numbers of cell divisions for cultured DN1 cells each day as measured by CTV. **D** Summary plots of the percentages of CD25-positive cells (blue) and percentage of Bcl11b-YFP-positive cells (red) from each day. The histograms show mean and standard deviation of data from two experiments.

Approximately 50% of DN1 cells turned on CD25 by day 2 of culture, with increasing percentages over time (Fig. 2B,D). However, few Bcl11b-YFP^+^ cells appeared among the CD25^+^ cells until day 3, with percentages of YFP^+^CD25^+^ cells then increasing over time (Fig. 2B,D). CTV analysis at day 2 showed that cell division was not required for CD25 expression (red dots=cells with 0 divisions), and that the most rapidly dividing cells were neither most or least likely to turn on CD25. By days 4-5, CD25^+^ cells had divided more rapidly than cells remaining CD25^−^, as seen by the shift towards a larger population with the lowest CTV levels (turquoise or lilac) (Fig. 2B). Bcl11b-YFP was not expressed in non-proliferating cells, and Bcl11b-YFP^+^ cells appeared to have proliferated more rapidly than cells remaining Bcl11b-YFP^−^ by days 4 and 5, since YFP^+^ cells generally retained lower CTV levels (Fig. 2B). Overall, these results confirm that CD25 is expressed before Bcl11b-YFP (Fig. 2B,D) and that CD25 can be turned on without cell division, while Bcl11b-YFP cannot. Furthermore, more developmentally advanced cells in these cultures proliferated somewhat faster than those that were delayed.

#### Clonal Kinetic Analysis of Differentiation from DN1

Because DN1s divide at different rates, the output populations become dominated over time by progeny of the most rapidly proliferating input cells. To resolve at a clonal level whether *Bcl11b* expression depends on time or cell cycles, individual thymic DN1 cells were co-cultured with OP9-DL1 stroma in microwells and microscopically imaged over time^24^ (Fig. 3A). DN1s were purified from thymi of mice with a constitutively-expressed membrane-(m)Tomato transgene, for cell segmentation and counts, as well as the nuclear Bcl11b-YFP reporter, and plated into microwells pre-seeded with OP9-DL1 cells. Microwells verified to start with single live DN1 cells were imaged daily for 6-7 days to determine clonal expansion and onsets of surface CD25 and nuclear Bcl11b-YFP expression within individual clones. Merged false-colour images of microwells in mTomato, CD25-AlexaFluor647, and Bcl11b-YFP fluorescence channels over time, are shown in Fig. 3B from one representative clone of 64 clones imaged. Two of the 64 DN1 clones generated only non-T cell lineage cells, possibly granulocytes, but the remainder turned on CD25 and most turned on Bcl11b-YFP within 7 days (Fig. S2). Fluorescence thresholds for determining CD25 and Bcl11b-YFP positivity were calculated using background fluorescence estimates (Fig. S3). Higher magnification images for cells from two representative clones from days 1-6 (Fig. 3C) show that at day 1 only the segmentation marker mTomato (grey) was expressed. In these clones, surface CD25 (magenta) was turned on by day 2 or 3 (wells 23 and 41 respectively), about 2 days before nuclear Bcl11b-YFP-positive cells (cyan) appeared by day 4 or 5.

**FIG. 3.**
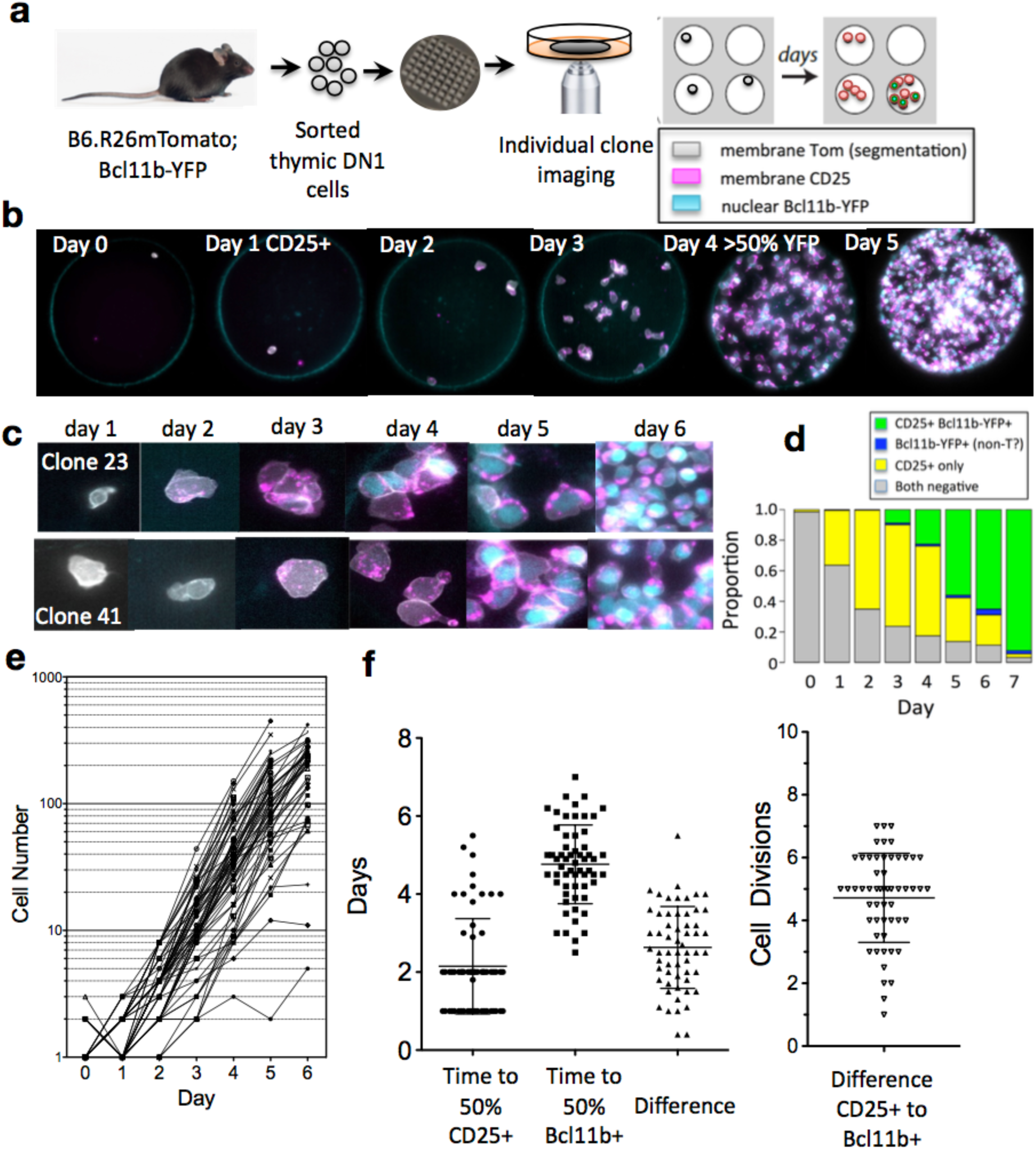
Kinetic analysis of proliferation and differentiation for individual DN1 clones. **A** Diagram of protocol used for imaging DN1 clones. Thymic DN1 cells were purified from mice homozygous for mTomato and the nuclear Bcl11b-YFP reporter, added to microwells pre-seeded with OP9-DL1 stroma, and cultured with cytokines IL-7 and Flt3L and CD25-AlexaFluor647 antibodies (marking progression to DN2). Individual microwells with one DN1 cell were imaged daily from day 0 to 6 or 7. **B** False-colour merged images of one DN1 clone for days 0 to 5: mTomato (gray), surface CD25 (magenta), and nuclear Bcl11b-YFP (cyan). Exposures were adjusted to allow visualization of cells in all channels. **C** Higher magnification merged false-colour images of selected cells from two DN1 clones showing the original DN1 cells lacking CD25 and Bcl11b-YFP expression on day 1 and the expression of CD25 (magenta) and Bcl11b-YFP (cyan) over time. **D** Combined proportions of cells expressing the two differentiation markers, CD25 and Bcl11b-YFP, alone or together over time. n=67 (including 62 T-lineage clones, 2 non-T clones, 3 non-clonal wells). **E** Log plots of cell numbers counted from daily images of 62 clonal T-lineage wells. Wells with only 1 cell on either day 1 or 2 were considered to be clonal. **F** (Left) Plot showing the timing of expression of CD25 and Bcl11b-YFP, scored as the days until >50% of cells in individual clones were positive for CD25 (n=62 clones) or Bcl11b-YFP (n=58), and the difference in time between the two events for the clones that turned on both markers during the experiment (n=58). (Right) Plot of the difference in cell cycles for individual clones between 50% of cells turning on CD25 and 50% turning on Bcl11b-YFP (n=58). Each symbol represents a single clone. Means and standard deviations are indicated with lines.

Differentiation dynamics for all imaged wells combined are shown in Fig. 3D. Most individual clones achieved similar proliferation rates but began with different lag times (Fig. 3E), reflecting heterogeneity within the starting DN1 population. At early timepoints, cells that remained DN1 had proliferated less than the CD25^+^ DN2s (Fig. S4), in agreement with CTV data. There was wide variability in the timing of CD25 and Bcl11b-YFP expression between clones (Figs. 3F, S2). CD25 came on most frequently on days 1-2 (76%) although 16% of clones did not turn on CD25 until >4-5 days of culture, while times to the onset of Bcl11b-YFP expression ranged from 3—7 days. CD25 up-regulation was not synchronous within some clones, hinting at a stochastic element even in the transition to DN2.

To test the determinism of commitment programming, we asked whether cells are programmed to turn on *Bcl11b* after a fixed period of time or fixed number of cell cycles after making the DN1-DN2 transition. Fig. 3F shows the differences for each clone between the times when >50% of cells were scored as CD25^+^ and as Bcl11b-YFP^+^, in elapsed time and in cell cycles (Fig. 3F). In fact, the intervals between entering the DN2 state (CD25^+^) and commitment (Bcl11b-YFP^+^) were highly variable both in absolute times and in cell cycles, taking from <1 to >5 days and from 1 to 7 cell divisions. Furthermore, Bcl11b-YFP was not turned on synchronously within a clone, generally requiring 2-3 days from expression in the first cells to expression in 100% of the cells (Figs. 3C (well 23, day 4), S2). Thus, daughter cells from individual DN1 cells are not only heterogeneous in their clonal founders’ differentiation states, but also show a large stochastic element in timing of *Bcl11b* up-regulation after entering the DN2 state^39^.

#### Single Cell RNA-FISH Measurement of Key Transcription Factors

Potential drivers of commitment include TCF1 (*Tcf7*), Gata3, and Runx1, which are necessary positive inputs for *Bcl11b* expression^24^, while PU.1 (*Spi1*) antagonizes developmental progression through commitment^23,40^. To determine the normal ranges of expression of *Spi1*, *Tcf7*, *Runx1* and *Gata3* in individual T-cell precursors in developmental stages leading up to *Bcl11b* activation for the model, DN thymocytes were purified from 5-week-old mice and analysed by multiplex single molecule RNA-FISH (smFISH) (Fig. 4A,B). Developmental stages of individual cells were scored based on measured Kit, CD25 protein and mRNA levels. Absolute transcript counts showed that *Runx1*, *Gata3*, and *Tcf7* expression was variable in DN1s, increasing up to 2x in DN2s (DN2 median transcript levels/cell: *Tcf7≍*40; *Runx1≍*12; *Gata3≍*8) (Fig. 4C). In contrast, *Spi1* was highest in DN1 and DN2, and declined with further development, in accord with previous bulk population measurements. *Bcl11b* was not expressed in DN1s but turned on in CD25^+^ DN2s (median transcripts/cell*≍*15), remaining high into DN3 (Fig. 4C).

**FIG. 4.**
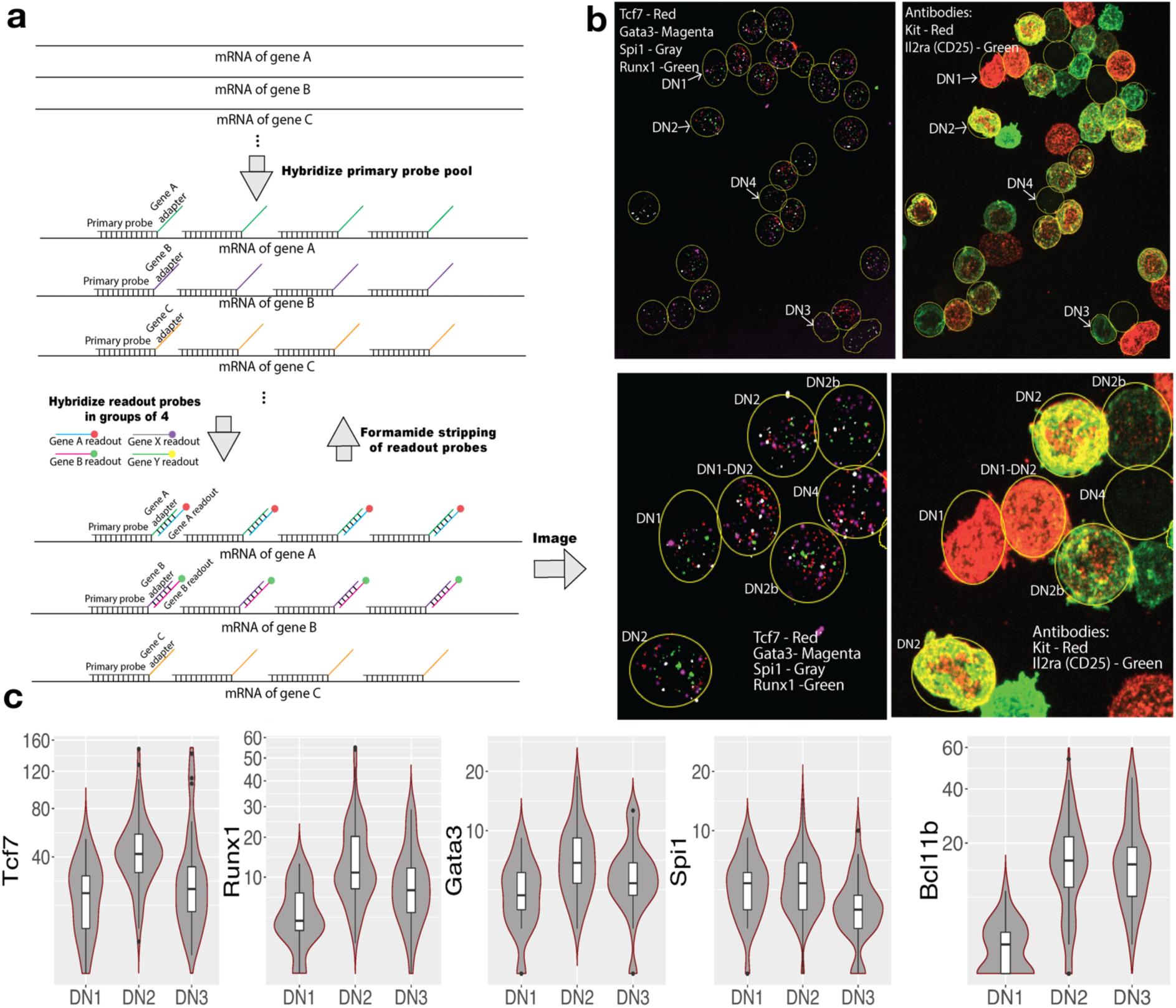
Regulatory gene expression measurement by smFISH. A. **A** Schematics of smFISH hybridizations. Primary probes were hybridized against mRNAs in immobilized DN cells in pools, and genes were detected with readout probes in groups of 4 genes, imaged and stripped with formamide before the second round of readout hybridizations. **B** Representative microscopy images. Each mRNA transcript was represented by a single dot under fluorescent microscopy, and the transcripts of each gene were counted (Tcf7, red; Gata3, magenta; Spi1, gray; Runx1, green) and assigned to individual cells that were categorized into DN1-DN2-DN3-DN4 by antibody stained surface markers Kit (red) and CD25 (encoded by Il2ra, green)). **C** Violin plots of transcript count distributions of *Runx1*, *Gata3*, *Spi1* (*PU.1*), *Tcf7* and *Bcl11b* in single cells, showing median and quartiles, in developmental stages DN1-DN3. n = 169.

These gene expression changes from DN1 to DN2 were used as input to the GRN model as an approximation for developmental changes in transcription factor protein. The half-lives of these proteins can differ substantially; for example, PU.1 has a half-life at least 10x that of TCF1, Runx1, and Gata3^36,41^, and so we used their relative expression patterns, rather than absolute protein levels.

### Modelling Results

As described earlier, a GRN model for *Bcl11b* activation during commitment should incorporate three separate aspects of control. First, four positive regulatory inputs, Notch signaling, TCF1, Gata3, and Runx1, are needed in the DN1-DN2a stages to make *Bcl11b* eligible for activation. Second, TCF1 and GATA3 work in a hit-and-run way and are dispensable for the final steps of *Bcl11b* induction, but Runx1 and Notch act through DN2a stage^24^. Third, activation of each *Bcl11b* allele, in a cell that has already fulfilled eligibility requirements, still requires a slow stochastic epigenetic remodelling process^39^. To accommodate these requirements, we separated an initial, deterministic process from a subsequent, stochastic process (Fig. 1B).

Thus, we developed a three-level dynamic model. (1) In the deterministic level, network dynamics between Notch signalling, Runx1, Tcf7, Gata3, and the antagonistic regulator PU.1 determine cell permissiveness for *Bcl11b* activation, via effects on the constraint function X. (2) In the stochastic level, the transition of *Bcl11b* from closed to open state is influenced by the inhibitory function, X, balanced against the continuing positive drivers Runx1 and Notch signalling. (3) In the population level, these single-cell processes are combined with a dynamic population growth model that accounts for the expansion of cells that have and have not yet activated *Bcl11b*. While the marker CD25 plays an important role in interpreting the experimental results relative to the DN1-DN2 transition, it is not a regulator and therefore is not explicitly included in these models. Model development and validation procedures are shown in Fig. 5.

**Fig. 5.**
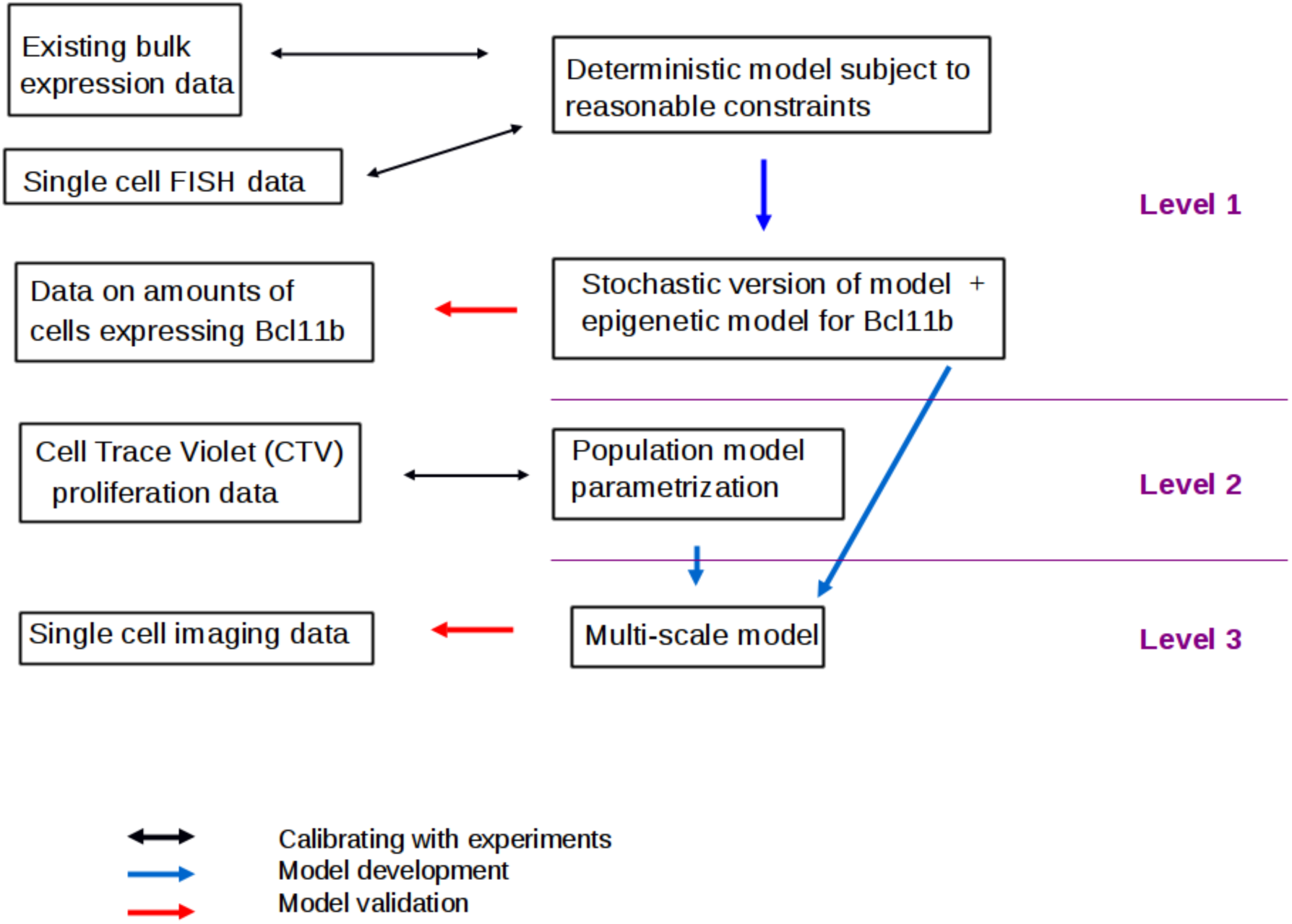
Flowchart of experimental and modelling steps emphasising predictions versus calibrations. **Level 1**: Bulk and single cell experiments are used to develop a dynamical model with both deterministic and stochastic versions. **Level 2**: The latter are augmented with an epigenetic model for *Bcl11b* expression. The resulting relative production rates of cells expressing *Bcl11b* are validated against bulk data. Cell Trace Violet data are then used to develop a population model for the different stages. **Level 3** This model is subsequently combined with the stochastic model from level 2 into a multi-level model that is validated against clonal imaging results.

#### The Gene Regulatory Network Architecture

For the first level of the single cell model we proposed a GRN architecture based upon gene expression, perturbation experiments and literature (Table 1), describing the interplay between the T-cell commitment genes *Tcf7*, *Gata3*, *Runx1* and the opposing gene *Spi1* (Fig. 6A). Rather than having these regulators controlling *Bcl11b* transcription directly, we proposed that their inputs collectively removed the obstruction from an antagonist of *Bcl11b* opening, X, representing a composite of the slow chromatin opening mechanism and the actions of any additional DN1-specific antagonists of *Bcl11b* expression, still to be defined. Only X, Notch signalling, and Runx1 then propagate to the next, stochastic level.

**FIG. 6.**
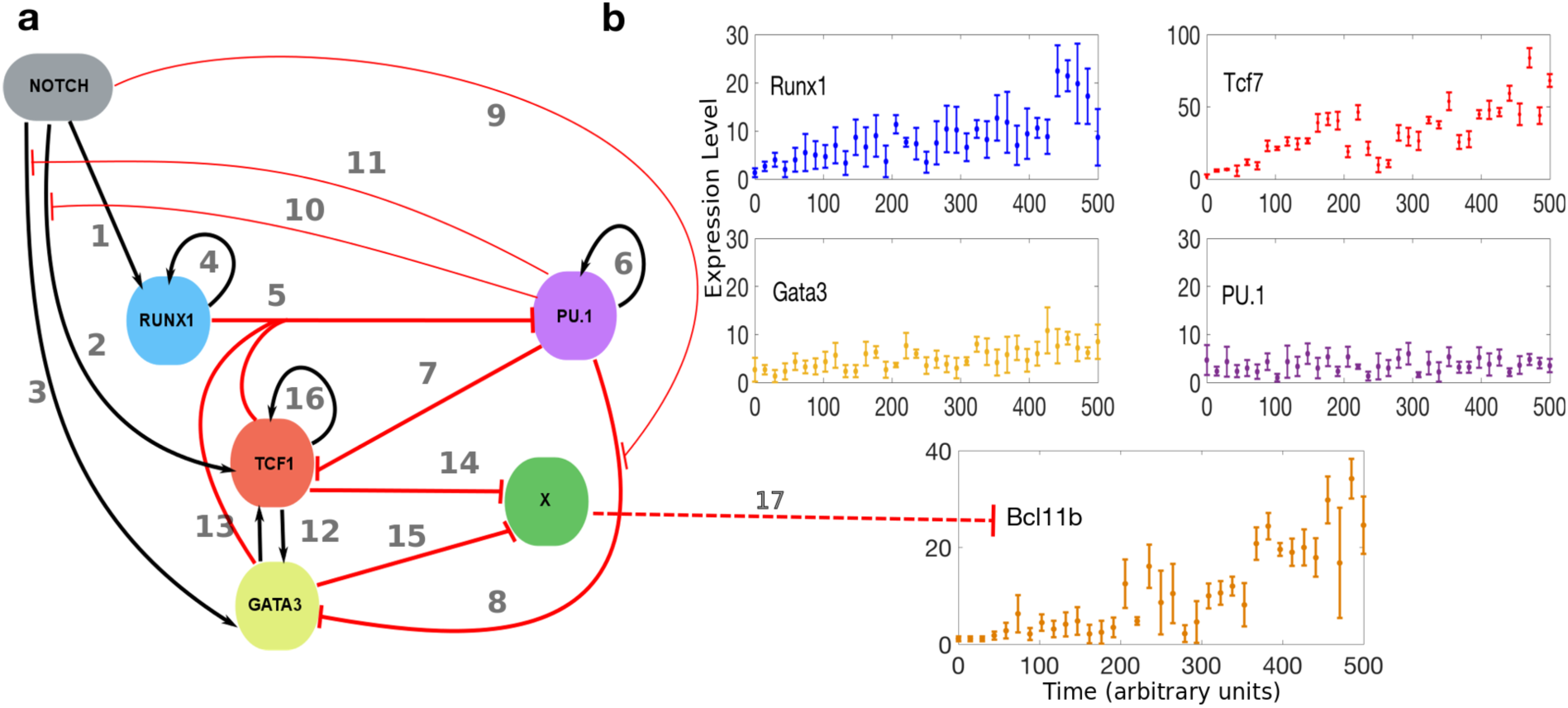
Core Circuit along with Pseudo-Time-Series data. **A** Gene Regulatory Network Topology, where each interaction has an index corresponding to the rightmost column in Table 1. The arrows indicate positive DNA regulation. The thick red lines with short bars represent repression of DNA regulation. The thin red lines with short bars depict PU.1 repression of Notch activation of Tcf1 and Gata3 and Notch inhibition of Pu.1 repression of Gata3 regulation. The dashed red line with short bar represents the function X causing epigenetic repression of Bcl11b. **B** Pseudo-Time-series obtained from ordering the clusters resulting from the Gaussian Mixture algorithm such as the resulting dynamics of Tcf7 and Bcl11b are similar to experimentally observed behaviour. Please note the different scales on the y axis, TCF7 having the highest counts therefore a scale to 100. The pseudo times-series data is shown as mean and standard deviation of each cluster, each dot depicting the mean value of the mRNA count for each cluster obtained from the FISH single molecule data

**TABLE 1:** Gene network components and sources of evidence.

| From | To | Effect | Evidence Type | References | Index |
| --- | --- | --- | --- | --- | --- |
| Notch | Runx1 | Activate | Modest up-regulation in pro-T cells | 21 | **1** |
| Notch | Tcf1 | Activate | Direct binding and activation | 16,17 | **2** |
| Notch | Gata3 | Activate | Developmental activation requirements and protection from repression, but no direct binding evidence | 8,53,56,57 | **3** |
| Runx1 | Runx1 | Activate | Auto-regulation seen in HSC precursors in embryo ('Cbfa2'); many binding sites; not certain in pro-T | 58 | **4** |
| Runx1 | PU.1 | Repress | Direct molecular evidence of repression through interaction sites | 11,15,26,59 | **5** |
| PU.1 | PU.1 | Activate | Functionally only in myeloid cells, promoter activation and auto-regulation via cell cycle | 6,18,41 | **6** |
| PU.1 | Tcf1 | Repress | Functional perturbation; possibly indirect due to paucity of binding | 10,21,23 | **7** |
| PU.1 | Gata3 | Repress | Functional perturbation; dependent on absence of Notch signal | 21 | **8** |
| Notch | Repression of Gata3 by PU.1 | Inhibit | Functional perturbation | 21 | **9** |
| PU.1 | Ability of Notch to activate Tcf1 | Inhibit | Functional perturbation | 10,21 | **10** |
| PU.1 | Ability of Notch to activate Gata3 | Inhibit | Functional perturbation | 21 | **11** |
| Tcf1 | Gata3 | Activate | Gain and loss of function perturbation in pro-T cells | 17 | **12** |
| Gata3 | Tcf1 | Activate | Genetic perturbation and retroviral interference | 19,22 | **13** |
| Tcf1 | X | Repress | Conjectural | Interpretation<br>24,39 | **14** |
| Gata3 | X | Repress | Conjectural | Interpretation<br>22,24,39 | **15** |
| Tcf1 | Tcf1 | Activate | Gain and loss of function perturbation in pro-T cells | 17 | **16** |
| X | Bcl11b | Repress | Epigenetic constraint on Bcl11b activation, apparently not relieved until after<br>DN1 to DN2 transition | 24,39 | **17** |

**TABLE 2.** Epigenetic model components and source of evidence.

| From | To | Effect | Evidence Type | References |
| --- | --- | --- | --- | --- |
| Runx1 | Bcl11b | Open | positively regulates *Bcl11b* expression amplitude at single-cell level based on gain and loss-of-function perturbation; direct DNA binding | 24,26,60 |
| Notch | Bcl11b | Open | accelerates and enhances frequency up-regulation from DN2; primes for activation in DN1 | 53, 24 |
| X | Bcl11b | Close | conjectural based on stage-dependent prohibition and inter-allelic asynchrony | 24, 39 |

#### Parameter Setting Based on Pseudo-Time-Series

From the single cell FISH data (Fig. 4), we created pseudo-time-series from DN1 through DN2, which show that in early DN1 expression of Tcf7 increases substantially and becomes predominant (Fig. 6B). After a delay (200 arbitrary time units), *Bcl11b* transcription begins. Furthermore, Gata3 expression slightly rises with increasing Tcf7 expression, along with Runx1. Expression of the opposing gene PU.1 is fairly constant at a low level through these stages, turning off later. The GRN model parameters (Table S1) were optimised from these pseudo-time-series (Fig. S5).

#### Transcriptional Level - Deterministic Simulations

We developed the computational model for the step 1 GRN (Fig. 6A) with Shea-Ackers deterministic rate equations^42^ (Methods). Deterministic simulation results from the model (with initial conditions for Tcf7, Gata3, PU.1 and Runx1 equal to the first pseudo-time-series expression values and the X function initially highly active) show that the system, when exposed to external Notch signalling, moves towards a steady state where the T-cell factors are highly expressed, with a pronounced increase in Tcf7, while PU.1 and X activity levels descend towards low values.

#### Transcriptional Level - Stochastic Simulations

Stochastic simulations^43^ of the GRN model (Fig. 7A) in individual cells with the same parameters (Table S1) show that noise could be a source of heterogeneity in DN1 cells, as not all simulations exhibited dynamic switching towards a steady state with high T-cell factors (Fig. 7C). Moreover, the stochastic simulations showed that GRN intrinsic noise sometimes leads to delays of the switch towards T-cell commitment.

**FIG. 7.**
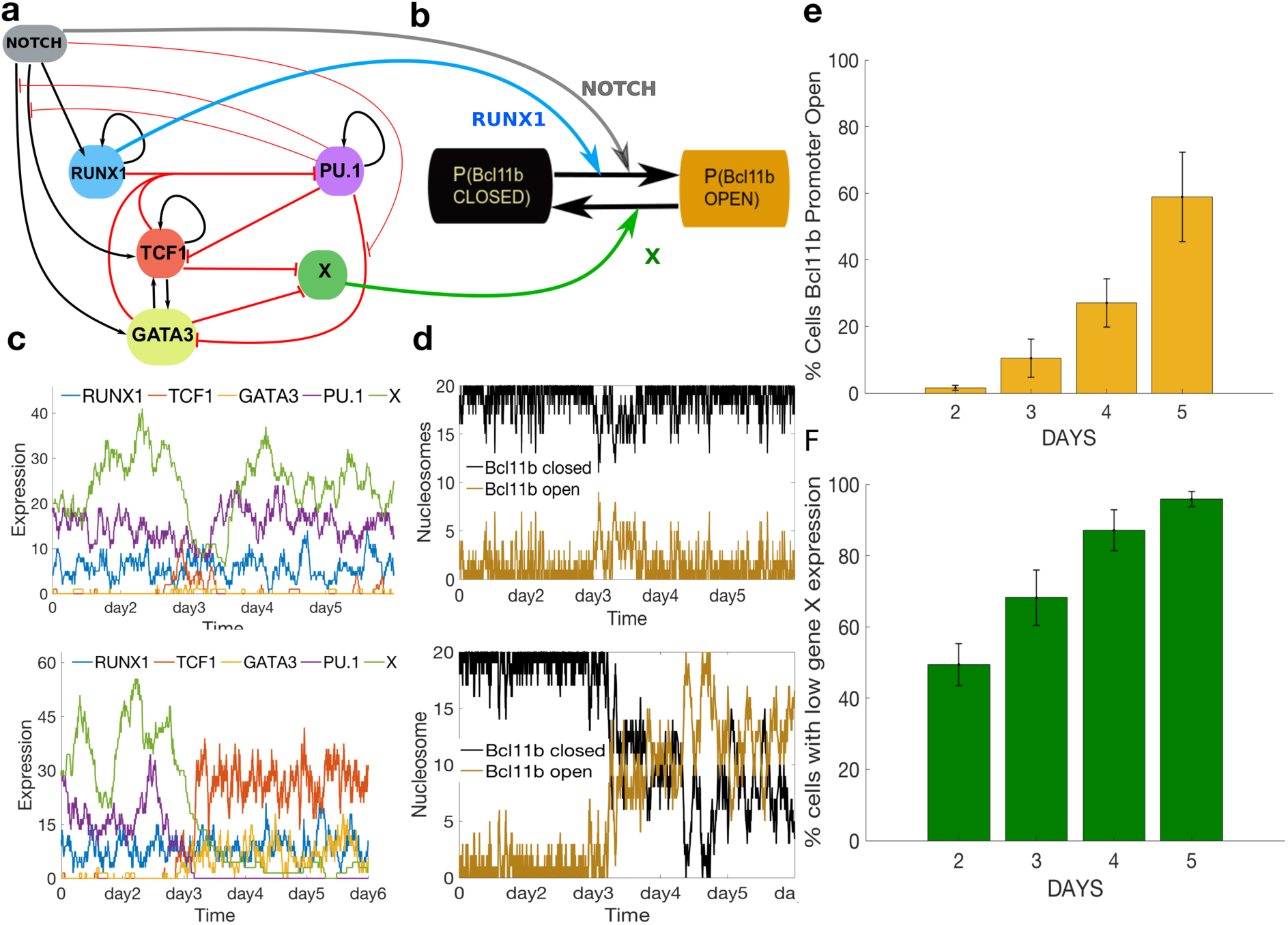
Single Cell Model Results. **A.** Gene regulatory network topology for transcriptional level of the single cell model where the thick red lines denote repression, the black arrows denote DNA regulation and the thin red lines denote inferred interactions from functional perturbation experiments. **B** Schematic representation of the epigenetic level model for the *Bcl11b* regulatory regions where Runx1 and Notch (blue and grey arrows) favour open DNA for transcription while X (green arrow) is keeping the *Bcl11b* regulatory regions closed. The green, blue and grey arrows show that X function activity levels along with Runx1 expression levels and Notch signalling activity serve as inputs for the epigenetic level of the single cell model. **C** Stochastic simulation results obtained from transcriptional level model. Simulation results where the system stays in a state controlled by Pu.1 and X are shown in the first time series plot. In the second plot, the T-cell factors are switched on while Pu.1 and X are down-regulated. **D** Stochastic simulations obtained from the epigenetic level model, shown in panel **B**, having inputs from the levels of Runx1, Notch and X obtained from stochastic simulations of the transcriptional level model. In the first plot, the multi-level single cell model stochastic simulation shows that the *Bcl11b* regulatory region stays closed while in the second plot the regulatory region becomes open. **E** Distribution (yellow) of the percentage of single-cell multi-level model simulations with the outcome of *Bcl11b* regulatory region open (mean and standard deviations for 3 sets of 100 single cells model simulations). **F** Distribution (green) of the percentage of single cell model simulations where X activity levels have low values at time points corresponding to each day of the experiment (mean and standard deviations for 3 sets of 100 single cells model simulations). The two distributions show that the model predicts a clear delay between the X loss of activity and *Bcl11b* regulatory region opening (e.g. at day 2 almost 50% of cells have lost X while less of 10% are *Bcl11b* positive).

#### Epigenetic Level

The second level of modelling consists of a simplified epigenetic model for the *Bcl11b* regulatory region with two possible states, open or closed, to account for the stochastic all-or-none activation timing observed for individual *Bcl11b* alleles (Fig. 8B). While the roles for Tcf7 and Gata3 apparently become dispensable immediately before *Bcl11b* activation, positive influences from Notch signalling and Runx1 remain. Notch signalling augments the likelihood of activation per unit time^24^, while Runx1 positively modulates expression levels^24,39^. Therefore, our model places these regulators in opposition to function X (Fig. 8A,B).

**FIG. 8.**
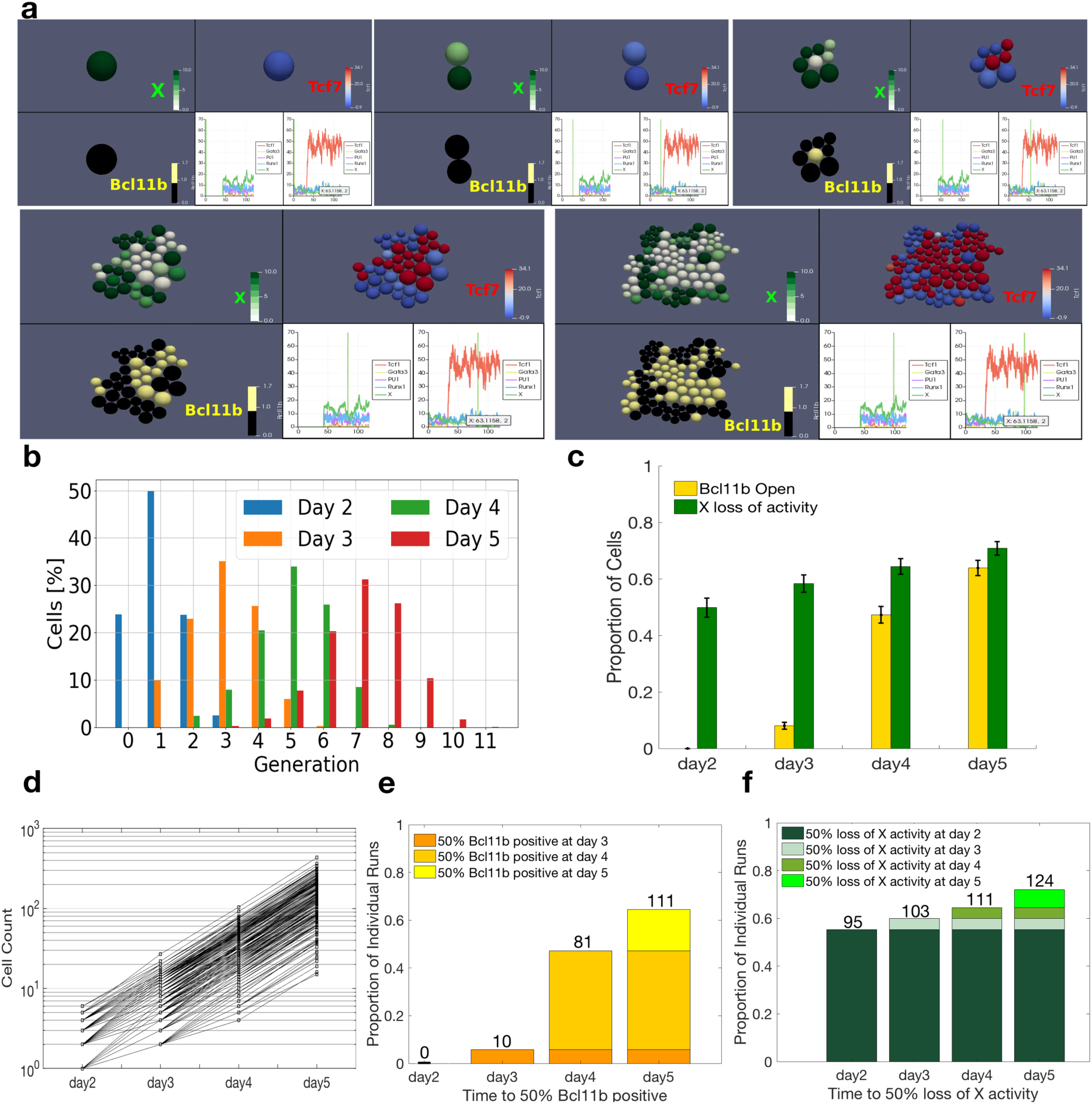
**A** Time snapshots of the evolving model cell culture starting from a single DN1 cell. Top left: Green scale shows X function activity impacting the epigenetic behaviour of the *Bcl11b* regulatory regions. Bottom left: Cells with opened/closed regions are indicated in gold/black respectively. Top right: Expression of *Tcf7*, which is predicted to be maximal during DN1-DN2a transition. Bottom right: Expression timeline of all genes for two selected cells; the first non-switching and the other switching to DN2a state. **B** Multi-scale model distributions of cell generations for each day. **C** Multi-scale model predictions of the mean of proportions of cells showing low X activity (green) along with the mean proportion of cells with the *Bcl11b* regulatory region open (yellow) at simulation times corresponding to day 2 to day 5 of CTV and single cell imaging experiments. Also shown are the standard errors from 172 simulations. **D** Number of cells from 172 simulations starting with one DN1 cell, showing the predictions on variability in proliferation kinetics between clones. **E** Multi-scale model predictions showing the proportion of simulations versus required time to reach 50% *Bcl11b* regulatory region open in each run. The number on top of the bins show the exact count of individual simulations reaching 50% at each day out of 172 runs. **E** Multi-scale model predictions showing the proportion of individual simulation runs versus the necessary time to reach 50% *Bcl11b* positive in each run. The numbers on top of the bins show the exact count of individual simulations reaching 50% at each day. **F** Multi-scale model predictions showing the proportion of individual simulations versus the time to reach 50% X activity loss among the simulated cells in each run. Also shown are the number of individual simulations reaching 50% at each day.

To account for the slow and potentially processive nature of this epigenetic transition, we used a model developed for calculating epigenetic activation as a function of the number of open (unmethylated) and closed (methylated) CpG dinucleotides at functional regulatory regions^44^ (Methods). Here it could apply to other probabilistic local chromatin states as well as DNA methylation. Notch signalling and Runx1 levels obtained from stochastic GRN simulations serve as positive regulators for the probabilities that CpG sites (or equivalent chromatin features that are propagated by similar chromatin “reader-writer” mechanisms) become open while X activity levels link to the probability of *Bcl11b* CpGs remaining closed.

#### Merging Transcriptional with Epigenetic Levels

We next conducted stochastic simulations of the single cell multi-level model incorporating *Bcl11b* regulatory region state dynamics predicting that early T-cell progenitor cells are heterogeneous in their CpG methylation states^37^ or equivalent open/closed chromatin states in the Bcl11b regulatory regions. The simulations showed that the *Bcl11b* locus can remain closed or become opened depending on the status of these CpG (or equivalent) sites, as dictated by Notch signalling, Runx1 expression, and opposing X activity (Fig. 7D). The stochastic treatment of the epigenetic level predicits further delays in T-cell commitment. In an example when the switch is thrown by day 3 (i.e., Tcf7 and Gata3 becoming highly expressed while *X* activity drops), the corresponding stochastic simulation of *Bcl11b* status showed that the regulatory system only opens after day 4 (Fig. 7C, lower panels).

Based on three sets of 100 stochastic simulations of this multi-level model, the model predicted that on days 2 and 3 <20% of cells would express *Bcl11b*, ∼40% on day 4, and ∼50% on day 5. These predictions agree well with direct monitoring of Bcl11b-YFP during 5 days in DN1-derived populations going through commitment (Fig. 2D, 3D). The distributions of cell percentages with low *X-*levels (Fig. 7F) are very similar to the distributions of CD25-positive cells (Fig. 2D, 3D), suggesting that loss of X activity might be correlated to CD25 expression, consistent with the observation that *Bcl11b* expression follows CD25 upregulation. Thus, the two-level single cell model shows that stochastic mechanisms controlling eligibility for chromatin opening downstream of the core deterministic GRN can account for the observed delays in T-cell progenitor commitment.

Furthermore, the model predicts the effects of knocking down *Tcf7, Gata3* and *Runx1*, which typically reach 15-25% of their original expression in RNA interference experiments. This corresponds to few molecular copies, thus requiring stochastic simulations. Early knock-down of *Tcf7, Gata3* or *Runx1* halts progression towards a state of high T-cell factors and low PU.1. Instead, the system results in a steady state with high PU.1 and low Tcf7, Gata3 and Runx1, keeping the *Bcl11b* locus closed (Fig. S6). In contrast, simulating the Tcf7 or Gata3 knock-down in DN2, after X is inactivated, showed no continued requirement for Tcf7 or Gata3 (Fig. S7), consistent with experimental findings^24^.

#### Population Model

To capture population dynamics, we developed population models based upon CTV data. We first tested the hypothesis that cell cycle length is independent of generation, which was clearly incorrect (Supplementary Material). We then relaxed this assumption, with each cell having its division time stochastically determined from a normal distribution with mean and variance depending upon cell generation (Fig. 8A). Normalised cell number distributions among different generations within simulated cell cultures up to day 5 were compared to the CTV data (Fig. 3C), allowing us to determine the cell cycle distribution parameters (Table S3). We monitored each cell generation, and calculated the proportion of cells in each generation at each day from 172 simulations (Fig. 8B) with resulting distributions very similar to experimental data (Fig. 3C).

#### Full Multi-scale Model

We next implemented the single-cell GRN model within the multi-cell simulation environment to determine how cell proliferation affects population distributions of T-cell fate-committed versus non-committed cells. To take into account the impact of cell division on the *Bcl11b* locus epigenetic state, we used an advanced version of the epigenetic model with an intermediate state between open and closed (Methods). Notably, the optimal number of “CpG equivalent” units that needed to be de-repressed was found to be 500, consistent with the very extended potential regulatory regions associated with the *Bcl11b* gene^32,37^.

We computed the proportion of cells for which the *Bcl11b* regulatory region was in the open state in 172 simulations starting with one cell proliferating for five days. The model predicts that none of the cells have the *Bcl11b* regulatory region open at day 2, while at day 3 roughly 8% are open. The proportion then increases to 47% at day 4 and 60% at day 5 (Fig. 8C). We also computed the percentage of cells that lose X activity at time points corresponding to each day (Fig. 8C). Our multi-scale model predicts that X activity is lost in ∼50% of the cells by day 2, and the proportion lacking X gradually increases, reaching >70% by day 5. This proportion of cells predicted to have lost X activity (Fig. 8C), interestingly, is close to the proportion of CD25-positive cells from experiments (Figs. 2D, 3D). Model predictions regarding the proportions of cells that lost X activity and then activated *Bcl11b* agree well with results from monitoring CD25 and Bcl11b-YFP expression in the developmental kinetics experiments (Fig. 2D, 3D).

Finally, the multi-scale model predictions were validated by imaging results from the 62 T-lineage DN1 clones (Fig. 3), corresponding to our model simulations starting with single clones and including cell divisions, GRN states and *Bcl11b* epigenetic dynamics. The single cell imaging experiments agree well with our model predictions that ∼50% of the clones contained ≥50% CD25^+^ cells at day 2 while almost none of the cells were yet Bcl11b-YFP-positive (Fig. 3D). The proportions of clones with ≥50% CD25^+^ and ≥50% *Bcl11*b^+^ cells then increased gradually, reaching more than 70% of clones by day 5 (Fig. 3D).

The multi-scale model also predicted the variability in proliferation kinetics between simulated single cell clones (Fig. 8D). The ‘in silico’ cell numbers were in good agreement with the observed ones (Fig. 3E). Furthermore, model predictions showed that the clones would not be fully synchronised, with 10/172 *in silico* “clones” producing 50% of *Bcl11b^+^* cells as early as day 3, with others waiting until day 4, 5 or even later (Fig. 8E). Similar results are obtained from monitoring the proportion of clones reaching 50% of cells without X activity (i.e., 95 of the simulated clones lost X activity at day 2 while 13 turned off X activity at day 5) (Fig. 8F). This agrees well with analysis for cultured and imaged DN1 clones, with variable timing for expression of both CD25 and *Bcl11b* (Fig. 3F). Thus, the predicted DN1 clonal heterogeneity in differentiation was validated by experimental data.

## DISCUSSION

We have developed, explored and validated an integrated computational multi-scale model for early T-cell development and commitment kinetics. Commitment of multipotent precursors to the T-cell fate is particularly accessible for modelling because, at the single-cell level, commitment corresponds to *Bcl11b* activation. Thus, the model explains commitment kinetics in terms of the gene network circuitry controlling *Bcl11b* expression by incorporating results from both single cell and population-level experiments elucidating the process on different scales. After a deterministic first level, our model is stochastic at the second level, including an epigenetic part to predict chromatin opening. Finally, the gene control model is integrated with a cell population model to align cell model properties with measured population behaviour.

Whereas other hematopoietic stem cell commitment processes have been subject to mathematical model investigations^34^, few attempts have been made for early T-cells^36,45^. Importantly, our model incorporates the complex molecular mechanisms shown recently to be involved in *Bcl11b* activation including the four known positive regulators contributing to the logic of *Bcl11b* activation, and the additional, slow epigenetic step that creates a stochastic delay in *Bcl11b* activation, long after the positive regulators are present. This level of control is probably not unique to *Bcl11b* regulation but has been demonstrated in particular depth for developmental activation of this gene^39^.

Computational models for epigenetics and GRNs were proposed in the context of pluripotency acquisition through cell reprogramming^44,46^. Multi-scale modelling approaches, unifying observations from the intra-cellular to cell population scale, have also been used to model heterogeneity and function of mature peripheral T-cells, reviewed in^47^. This is, to our knowledge, the first time a computational model, which considers the epigenetic and genetic levels of regulation, is used to explain and predict dynamics of T-cell lineage commitment.

The choice to use continuous-valued rather than Boolean transcriptional models has the advantage of avoiding discretising the data in instances when experimental real-valued gene expression levels are relevant for the problem studied, as in this case. On the other hand, with a Boolean approach one can probe larger networks than that modeled here. These issues have been briefly reviewed^34^. We have chosen the continuous-valued approach as our current data provide information beyond on/off representations, and previous investigations of important regulators of *Bcl11b* enable us to focus on a few key genes.

The design of the stochastic epigenetic level of the network was novel, incorporating some features not usually included in GRNs, in order to encompass the known mechanisms involved. Published experimental measurements indicate that a default repressive chromatin state is probably a major contributor to the stochastic timing of activation, with a delay from the activation of one allele to the other - a measure of noise in time to relieve an epigenetic barrier - on the order of days^39^. We linked relief of this negative epigenetic function (X) to the observed developmental timing control, because *Bcl11b* is only turned on in the DN2a stage, never (or rarely) in DN1, even though all the known positive regulators are already present in DN1. For parsimony, we have proposed an architecture in which the positive regulators in the first deterministic layer of the model work toward down-regulating X, leaving the second stochastic layer to determine when *Bcl11b* will respond by activation.

Importantly, X is a composite function in this model, not a single undefined regulatory gene. As *Bcl11b* starts out in an epigenetically silent state inherited from hematopoietic stem cells, X measures the resistance to the opening of relevant cis-regulatory elements at each allele of the locus. It is possible that some transcription factors inherited from stem/multipotent progenitors and still active in DN1 could sustain X, but this epigenetic constraint itself cannot be equated with a trans-acting repressor. Therefore, in the stochastic level of the model, we have taken the balance of Notch signals and Runx1 against decreasing X to drive a stepwise epigenetic opening process that we have modelled analogously to the removal of methylated CpGs. While demethylation may be part of this process, the use of this formalism is more abstract here and indicates the stochastic but neighbour-biased and progressive nature that is shared by demethylation with other epigenetic opening processes.

Stochastic simulations of the GRN controlling the first steps in T-cell commitment show that the switch towards a state with low X-levels does not occur in a synchronised manner between model simulations with identical initial conditions. This suggests that GRN intrinsic noise could be a source of heterogeneity in the loss of X, which is one of the first steps in the T-cell commitment process and we speculate that it could be linked to the DN1 to DN2 transition marked by CD25 expression.

CD25 expression, the key marker of transition to DN2, was monitored in experiments but not included in the model. Nevertheless it provides several key insights into *Bcl11b* expression kinetics. First, the timing of CD25 expression is quite heterogeneous in different DN1 clones, confirming DN1 cell heterogeneity^1,35^. Furthermore, the clonal analysis shows that CD25^+^ DN2 cells divide more quickly than CD25^−^ DN1s. One ramification of more rapid cell division is the dilution of proteins affecting the potential speed of epigenetic changes^48^. Finally, both CTV and clonal imaging confirm that *Bcl11b* is typically turned on days and many cell cycles after CD25, in every clone (Fig. 2,3). This is surprisingly comparable to the average and range of delays predicted between X function down-regulation and *Bcl11b* opening in the model, hinting at possible common regulatory elements yet to be determined.

In summary, this new model that integrates three distinct types of mechanistic control provides a quantitative understanding of commitment kinetics in early T cell precursors, and a valuable *in silico* tool to help discover new commitment regulators in this system.

## METHODS

### Mice

C57Bl/6, B6.Bcl11b^yfp/yfp^ (Bcl11btm1.1Evr) reporter mice^24^, and B6.129(Cg)-Gt(ROSA)26Sor^{tm4(ACTB-tdTomato,-EGFP)^ mice expressing ubiquitous membrane Tomato (Jackson Laboratory) were bred and maintained in the Caltech Lab Animal Resources Facility in accordance with protocols reviewed and approved by the Animal Care and Use Committee at California Institute of Technology. Note that the *Bcl11b^yfp^* allele is a nondisruptive insertion of an internal ribosome entry site (IRES)-mCitrine (YFP) reporter into the 3’ untranslated region of the last *Bcl11b* exon, which allows fully normal expression of the Bcl11b protein from the same allele.

### DN cell purification

Single cell suspensions were made from thymuses from 4-6-week-old mice. Thymuses were removed, passed through sterile metal meshes and collected in 1X Hanks Balanced Salt Solution supplemented with 0.5 % fraction V bovine serum albumin, 10mM Hepes buffer (Gibco), 5mM MgCl2, and DNAseI. After pelleting, the cells were resuspended in a cocktail of biotinylated antibodies to deplete unwanted cells: CD8a (53-6.7), TCRgd (GL3), TCRb (572597), Gr1 (R86.8C5), Ter119 (Ter119), NK1.1 (PK136), CD11b, and CD11c (N418), after which the cells were incubated with streptavidin-coated magnetic beads and then passed through a magnetic column (Miltenyi Biotec). The eluted DN cells were either used directly for RNA-FISH analysis or further purified by staining with fluorescent antibodies to CD44, Kit, CD25 and then sorting on a BD Biosciences FACSAria or FACSAriaFusion or a Sony SY3200 Cell Sorter in the Caltech Flow Cytometry Facility. For bulk cultures or clonal imaging DN1 cells were sorted as CD44high Kit^high^ CD25-negative Bcl11b-YFP-negative cells. Note that for simplicity, throughout this paper, we use “DN1” to refer to the Kit^high^ subset of CD44+ CD25-cells that contains the T-cell precursor function, also known as ETP.

### Single Molecule Multiplex Fluorescent in situ Hybridization (sm-FISH)

#### Probe design and synthesis

The gene-specific primary probes for each gene tested were designed as previously described{Shah, 2016 #2362}, with some modifications: each probe comprised a mRNA-complementary 35-mer sequence plus a shorter “gene handle” sequence that was shared by all probes against the same gene. All probes were blasted against the mouse transcriptome and expected copy numbers of off-target probe hits were calculated using predicted RNA counts in the ENCODE database for murine thymocytes. BLAST hits on any sequences other than the target gene with a 15-nt match were considered off-target hits. Any probe that hit an expected total off-target copy number exceeding 500 in count table was dropped. Probes were sequentially dropped from genes until any off-target gene was hit by no more than 6 probes from the entire pool. At this stage, all of the viable candidate probes for each gene had been identified. For the final probe set, the best possible subset from the viable probes for each gene was selected such that none of the final probes used were within 2-nt bases of each other on the target mRNA sequence, with no-overlapping hybridization regions and GC-content close to 55%. Primary probes were synthesised and amplified from array-synthesised oligo-pool as previously described^49^. The readout oligos against specific gene handles on primary probes were ordered from IDT (Integrated DNA Technologies, Coralville, Iowa) with 5’-amino modification and were coupled with NHS (N-hydroxysuccinimide)-ester labelled fluorescent dyes (Alexa 488, 594, 647 (Thermo Fisher Scientific) and Cy3B (GE Healthcare)) and purified through HPLC.

#### smFISH experiment and image acquisition

First, the isolated cells were spun onto an aminosilane modified coverslip, crosslinked with 4 % Formaldehyde (ThermoScientific 28908) in 1X PBS for 10min, and permeabilized in 70 % EtOH overnight at 4C. Samples were imaged first to record the surface antibody signals, followed by briefly bleaching away antibody signals through incubation in 0.1 % NaBH_4_ (Sigma 452882) in 1XPBS for 10 min. Then, the samples were **1)** hybridised overnight at 37° C with primary mRNA probes at 1 nM each oligo concentration in 50% Hybridization Buffer (50% HB: 2X SSC (saline sodium citrate, Invitrogen 15557-036), 50% (v/v) Formamide (Ambion AM9344), 10 % Dextran Sulfate (Sigma D8906) in Ultrapure water (Invitrogen 10977-015)); then **2)** washed in 50 % Wash Buffer [2X SSC, 50 % (v/v) Formamide, 0.1 % Triton-X 100 (Sigma X-100)] for 20 minutes, followed by incubation in 2X SSC for 10 minutes. The samples were then **3)** incubated with fluorophore-coupled readout oligos, in 30 % Hybridization buffer (30% HB: 2X SSC, 30% (v/v) Formaldehyde, 10% Dextran Sulfate in Ultrapure water) at concentrations of 10 nM each oligonucleotide for 30 minutes; this was followed by **4)** 5 minutes wash in 30 % Wash Buffer (2X SSC, 30 % Formamide (v/v), 0.1 % Triton-X 100 (Sigma X-100)), 3 minute wash in 2X SSC and DAPI staining. We then **5)** proceeded to imaging of this round of hybridization as described below. After image acquisition, 6) the samples were incubated with 70% formamide with 1x PBS at room temperature for 30 minutes, followed by 3 rounds of washing in 1x PBS for 5 minutes each round. The procedures **3)-6)** were then repeated with a set of gene-specific readout oligos until the completion of measurements of all genes of interest, as illustrated in Fig. 4A.

Samples were imaged in an anti-bleaching buffer (20 mM Tris-HCl, 50 mM NaCl, 0.8 % glucose, saturated Trolox (Acros Organics 218940050), pyranose oxidase (OD405 = 0.05) (Sigma P4234), and catalase at a dilution of 1/1000 (Sigma C3155)) with the microscope (Olympus IX81) equipped with a confocal scanner unit (Yokogawa CSU-W1), a CCD camera (Andor iKon-M 934), 100x oil objective lens (Olympus NA 1.4), and a motorised stage (ASI MS2000). Lasers from CNI and filter sets from Semrock were used. Snapshots were acquired with 0.5 μm z steps for more than 10 positions per sample.

The cells were segmented and categorised according to surface antibodies, and dots representing individual mRNA molecules were assigned to individual cells as shown in Fig. 4B.

### Bulk cell cultures

For experiments on developmental kinetics DN1 cells were cultured on OP9-DL1 stromal cells as previously described^24,29^ in *α*MEM medium supplemented with L-glutamine, penicillin, streptomycin (OP9 culture medium) and IL-7 (5 ng/ml) and Flt3L (10 ng/ml). For tracking proliferation in bulk cultures, FACS purified DN1 cells were stained in 5 μM Cell Trace Violet (Molecular Probes) for 7 minutes at 37°C in HBSS and washed 2X with whole medium before being plated in 96-well OP9-DL1 co-cultures. Cells were incubated at 37°C 7% CO_2_, harvested on days 2, 3, 4, and 5 by vigorous pipetting, and evaluated for developmental status by staining with antibodies, CD25 and Kit for development and CD45 to separate developing cells from stroma, plus a viability dye, 7AAD. The cells were then analysed for expression of these markers plus Bcl11b-YFP using a Miltenyi MACSQUANT flow cytometer. Analysis was carried out using FlowJo software (Treestar). CTV measurements of cell division histories were calibrated using total DN (DN1—4) cultures on OP9-DL1 stroma (Fig. S1).

### Clonal Image Analysis

To follow the kinetics of development and proliferation in individual DN1-derived clones by microscopic imaging, several technical problems had to be solved. First, to allow for accurate identification and segmentation of developing cells of varying shapes and sizes on a background layer of OP9-DL1 stroma, we needed to use cells purified from B6.R26m^mTom/mTom^ Bcl11b^yfp/yfp^ J mice expressing ubiquitous membrane Tomato (Jackson Laboratory). We then crossed and backcrossed this strain to our B6.Bcl11b^yfp/yfp^ reporter mice ^24^ until both loci were homozygous (B6.ROSA26-mTom;Bcl11b-YFP mice). To enable the use of the Bcl11b-YFP reporter for imaging the commitment status of individual developing T-cell precursors, GFP had to be removed from the original OP9-DL1 stromal cells^50^. To do this, we designed guide RNAs targeting GFP and electroporated them along with a CRISPR puromycin plasmid v2.0 (Plasmid # 62988, Addgene) into OP9-DL1-GFP cells using the Lonza Nucleofector Kit. After puromycin selection, different clones were analysed for GFP expression and one clone found to be completely negative for GFP (OP9-DL1-delGFP1) which was selected for use in imaging.

Developing T-cells are extremely active in motility, so individual DN1 cells had to be confined in wells for imaging. We used a protocol similar to our previous experiments with DN2 cells ^24^, but we found that the background of the clear wells was too high in the fluorescence channels used in these experiments, so black microwells were substituted. Black 8mm circular poly (dimethyl siloxane) PDMS micromeshes with over 150 microwells punched/micromesh, each approximately 250 μM wide X 100 μM deep, were custom fabricated by Microsurfaces (Australia). One day before the start of co-culture and imaging these micro meshes were adhered to 2-4 (macro)wells of a 24-well glass-bottom plate (P24G-1.0-13-F) (MatTek, Ashland, CA), sterilized with ethanol, and washed in accordance with manufacturer’s instructions, and then seeded with 5,000 OP9-DL1-delGFP stromal cells. Cultures were carried out in OP9 culture medium prepared as previously described except for the omission of pH indicator, phenol red, from the medium, which gives background fluorescence, and with the addition of 10mM Hepes buffer to stabilize the pH of the wells during imaging. On the first day of culture the following were added to the medium: 10 ng/ml Flt3L, 5 ng/ml IL-7, and 0.05 μg/ml anti-CD25-AlexaFluor647 antibody (BioLegend) for detection of CD25 surface expression. DN cells were isolated from B6.ROSA26-mTom/Bcl11b-YFP mice and DN1 cells were FACS-purified as described above. 1,500 sorted DN1 cells were added to each well of the 24-well plate which had the microwells pre-seeded with OP9-DL1-delGFP. The cells were allowed to recover for at least one-hour at 37C, 7% CO_2_, before transfer to the Leica 6000 wide-field fluorescence microscope with Metamorph software and an incubation chamber pre-set to 37C, 7 % CO_2_. Each microwell was briefly checked using the 40X objective, which allowed imaging of one entire microwell, for mTomato-positive cells. Microwells with 1 or 2 cells had their X-Y stage positions marked. All marked microwells were imaged daily using differential interference contrast (DIC) and three fluorescence channels: 504 excitation-542 filter for YFP, 560-607 for mTomato, and 650-684 for CD25-AlexaFluor647. Wells found to have one mTomato positive cell on either day 1 or 2 and cells present throughout the culture period were selected to further analysis. mTomato-positive cells were segmented and analysed using Fiji (ImageJ) and by hand to obtain cell area and CD25 and Bcl11b-YFP fluorescence data. For each cell, total fluorescence was calculated by multiplying mean fluorescence by area. Because microwells became quite crowded by late time points (days 6-7), and cell clumps were very difficult to segment, all cells were counted but only a subset of the distinguishable cells were segmented and sampled to obtain fluorescence data for the well. Fifty background fluorescence sample cells were taken and the mean total fluorescence for CD25 and Bcl11b-YFP plus 3 standard deviations was used to set the thresholds for determination of positive vs. negative cells.

### Pseudo time series

From the single cell FISH data, pseudo-time-series were created by clustering the data and ordering the clusters, assuming that the cells measured are in states distributed over the whole-time intervals of both stages DN1 and DN2. The data points were first classified into clusters with a Gaussian-Mixture algorithm^51^, assuming that they follow several Gaussian distributions with individual parameters. Moreover, the number of data points in each cluster should be similar in order to regard all data with the same significance. The mean expression and standard deviation of all data points in a cluster were then computed for each gene and used to arrange the clusters in time. The cluster order was set by the fact that both *Tcf7* and *Bcl11b* expression levels are known to increase during cell commitment^52,53^. We considered the relative values of gene expression levels with respect to the maximum level at each stage to prohibit Tcf7 from dictating the order of the clusters alone, since its absolute expression is higher than *Bcl11b* expression. For each cluster, expression values of *Tcf7* and *Bcl11b* were added. Finally, the clusters were ordered such that this sum is increasing and placed equidistantly on an arbitrary time axis.

### Single Cell Model - Transcription Level

For the circuit in Fig. 6A we obtained the following set of rate equations from a thermodynamic approach^42^. The equations describe the dynamics of T-cell specific genes *Tcf7*, *Gata3* and *Runx1* along with gene opposing the T-cell fate *PU.1* and X function, with concentration levels denoted as: [T], [G], [R], [P] and [X]. The Notch signalling activity is denoted as N.

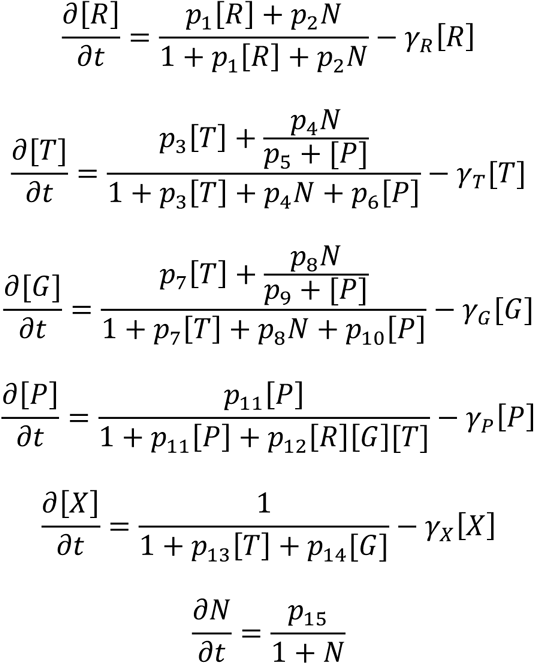

We considered that repression of Pu.1 by the T-cell specific factors follows AND logic i.e. Runx1, Tcf7 and Gata3 all need to be present for active repression of Pu.1. This accounts for the experimentally observed increases in Pu.1 expression upon deletion or downregulation of any one of these regulators (see Table 1 for references). We modelled the Notch activity such that the N levels increase throughout the simulations, however, parameter p_15_ values were chosen to be very low to assure only a slight increase of the Notch activity, avoiding a jump to infinity. Pu.1 limits the up-regulation of Tcf7 and Gata3 by Notch signalling. This was modelled by dividing the Notch activation term, in the nominator of the rate equations of Tcf7 and Gata3, by a linear term including Pu.1 expression level. The rest of interactions in the model follow a standard Shea-Ackers formalism implementation. The parameters p_i_ with i=1:14 except p_5_, p_9_, which are part of implementation of Pu.1 impeding the Notch positive regulation of Tcf7 and Gata3, correspond to binding affinities, while *γ* parameters model the decay rates linked to the half-lives of the respective molecules. The parameter values (Table S1) were chosen such as the resulting expression levels dynamics for all the genes in the network are within the range levels of genes expression observed in the single cell FISH experiments (Fig. S6). The model parameters were optimised (Supplementary Material) from the pseudo-time-series data using simulated annealing, genetic algorithms and a bound constrained optimisation algorithm (L-BFGS-B)^54^ with the latter performing best.

### Single Cell Model - Epigenetic Level

We implemented a simplified computational model for *Bcl11b* regulatory system transition from chromatin closed to open state. The epigenetic level model is linked to the transcription level model through the fact that the expression of *Runx1* and X and level of activity of Notch signalling (outputs of transcription level model) serve as inputs to the *Bcl11b* regulatory system model. We consider that Runx1 along with Notch signalling activity push towards an open *Bcl11b* regulatory system state thus the probability of achieving this state is direct proportional to Runx1 and Notch levels. X activity is considered to help maintain a closed state of the *Bcl11b* regulatory system, dictating the probability of the regulatory region to be closed. We initially put forward a very simplified model in which we considered the size of the Bcl11b regulatory system to have a finite low number of CpG sites i.e. 20. This was chosen to make sure that the simulations were very fast, but that we considered enough CpG sites so that the transitions between the opened and closed states are not instant and not completely controlled by noise. The amount of existing CpG sites in one of the states influences negatively the amount of CpG sites in the other state, because of the finite total number of CpG sites. We conducted a stochastic implementation of the simplified model for epigenetic state evolution of Bcl11b regulatory region governed by the following master equation:

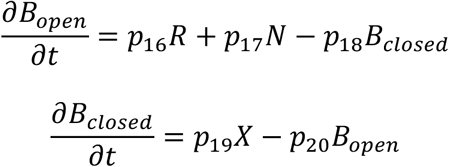

where N denotes Notch signalling activity, R represents the expression level of Runx1 and X is the X activity level. The model parameter values are shown in Table S2.

The initial conditions for stochastic simulations of the epigenetic level model consider the *Bcl11b* regulatory system to be in closed state i.e. the number of closed CpG sites is greater than the number of sites considered to be open. In some simulations the *Bcl11b* regulatory system becomes open i.e. the number of opened CpG sites becomes and remains greater than the number of closed CpG sites during simulation time.

### Multi-scale model of proliferation and gene expression

To analyse the effects of cell proliferation on the population scale gene expression we devised a model in which we can track the gene expression and division history of each individual cell. The model is an extension of in house developed framework, written in C++ and used previously in the context of cell-based simulations^55^. The mechanical interactions between cells are modelled in the same manner as in the original implementation. Each cell in the model contains a copy of a gene network presented in Single Cell Model (both transcription and epigenetic levels) and evolves its gene expression levels independently from other cells by a stochastic Gillespie simulation^43^. Cell cycle length for each cell is also a stochastic variable and chosen from a normal distribution. The cell divisions are assumed to be symmetric such that the daughter cells inherit the mother cell content, but then evolve independently. Since we assume that the presented network describes transition from DN1 to DN2a cell state in the multi-cell simulations we turn on the network in its initial state between day 1 and day 2 while the start of the simulation at day 0 corresponds to introduction of a single DN1 clone to the well.

We constructed a unified model of cell proliferation consistent with the data by assuming that the cell cycle length is a function of the cell generation. The parameters of the cell cycle length normal distributions were chosen independently for each cell based on its generation and fitted globally to the CTV data in Table S4.

In order to take into account the effect of cell division on the amounts of epigenetic factors inside a cell, we implemented a Bcl11b regulation region epigenetic model where the regulatory system can be open, close or at an intermediate state corresponding, in a CpG site methylation model, to the methylated, unmethylated and hemi-methylated states (Fig. S8). This type of collaborative model was proposed in^44,48^ and used here to simulate the opening and closing of the *Bcl11b* regulation region when cells go under division process. In the model the CpG sites can be methylated (M) -- closed state, hemi-methylated (H) -- intermediate state or unmethylated (U) -- open state. Transition rate constants depend on the level of X activity, expression level of Runx1 as well as Notch signalling activity. The arrows labelled with *α* to *ε* indicate reactions that require a mediator nearby (e.g. the *α* arrow defines a transition from U to H in the presence of a mediator in state M).

The collaborative model is described be the following set of equations:

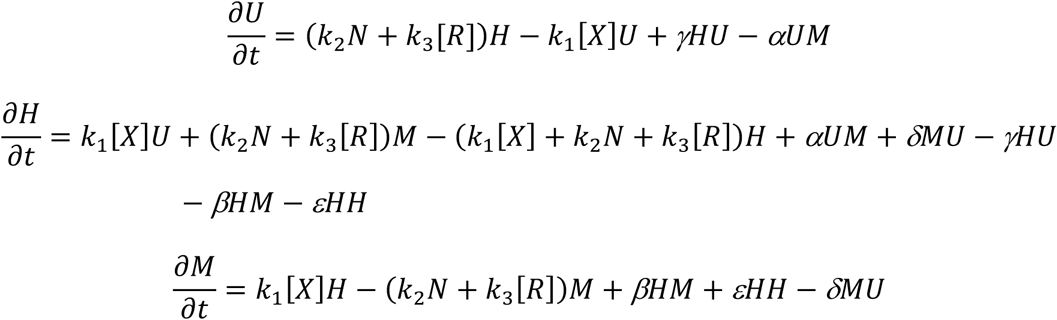

The parameter values are shown in Table S5.

When we simulate division in the multi-scale model, two new Gillespie simulations are started. In the absence of X, and in response to Runx1 (R) and Notch signals (N), the states of the mother cell *Bcl11b* regulatory region CpG sites are transferred to the daughters following the rule: M -> H, H -> 50% H + 50% U, U -> U^48^. It should be noted that when the collaborative model was used, the *Bcl11b* regulatory system had to be of a size of minimum 500 CpG sites. The results shown in Fig. 8C were obtained only if the regulatory region of *Bcl11b* went through epigenetic events corresponding to demethylation of at least 500 CpG sites. Note that the real *Bcl11b* gene includes multiple methylated CpG islands in the gene body and also possesses extended enhancers spread over more than 850 million bp of DNA^32,37^, so this constraint is quite likely to be biologically meaningful.

## ACKNOWLEDGEMENTS

The authors thank Dr. Long Cai for support for the smFISH analysis; Dr. Jeffrey Longmate for data analysis; Dr. Hao Yuan Kueh for helpful discussions and advice on imaging and analysis; Kenneth Ng for technical help; Diana Perez, Jamie Tijerina, and Rochelle Diamond of the Caltech Flow Cytometry Facility for FACS cell sorting; Dr. Andreas Collazo and the Caltech Biological Imaging Facility for microscopy assistance. The authors gratefully acknowledge the support of the US National Institutes of Health (USPHS grant R01HL119102 to E.V.R. and C.P.) and the Albert Billings Ruddock Professorship (to E.V.R)

## AUTHOR CONTRIBUTIONS

VO, MY, EVR and CP designed the study.

VO, MY, EVR and CP wrote most of the manuscript.

MY performed the CTV and kinetics experiments.

WZ performed the FISH experiments.

VO developed the transcriptional and epigenetic models and analyzed the data.

PK developed the population model.

JD implemented the pseudo time series and multi-scale models.

