## Supplementary Material for "Multi-scale dynamical modelling of T-cell development from an early thymic progenitor state to lineage commitment"

### *Supplementary Information*

Victor Olariu[1†], Mary A. Yui[2†], Pawel Krupinski[1], Wen Zhou[2], Julia Deichmann[1], Ellen V. Rothenberg[2*], Carsten Peterson[1*]

[1] Computational Biology and Biological Physics, Lund University, Lund, Sweden

[2] Division of Biology and Biological Engineering, 156-29, California Institute of Technology, Pasadena, California 91125, USA

[†] These authors contributed equally to this work

#### Parameter Optimization - Single Cell Model

For the transcriptional level of the single cell model we used a cost function of the model parameter set **p** measuring the squared difference between the pseudo-time-series data D and model simulated concentration values of the genes in the network, all grouped under variable M:

$$S(\boldsymbol{p}) = \sum_{k}\sum_{i=1}^{n}(D_{k,j} - M_{k,j}(\boldsymbol{p}))^2 \; where \; k = [R],[T],[G],[P],N$$

n is the number of data points for each gene in the pseudo-time-series data, genes *Tcf7*, *Gata3* and *Runx1* along with gene opposing the T-cell fate *PU.1* and X function, concentration levels were denoted as: [T], [G], [R], [P] and [X]. The Notch signalling activity is denoted as N.

S(**p**) was minimized with respect to the model parameters using multiple algorithms e.g. simulated annealing, genetic algorithms and a bound constrained optimisation algorithm (L-BFGS-B)[57]. The latter performed best.

The deterministic and stochastic simulation results presented in Fig. 8 were obtained using the parameter values shown in Table 4. Among the parameters, the decay rates are given in real units, whereas the other parameters cannot be tied with experimental numbers as there are overall implicit constants multiplying the ratios appearing on the right-hand side of the rate equations.

It should be noted that we consider the starting state of the system to correspond to an uncommitted early progenitor cell with high expression levels of *PU.1* and X activity along with *Runx1* being expressed and low expression levels of *Tcf7*, *Gata3*. In actual DN1-DN2a stage cells, note that several other progenitor-specific transcription factors in addition to *PU.1* are still expressed until commitment, but the network connections of these factors to *Tcf7*, *Gata3*, *Runx1* and Notch signalling have not yet been studied in depth 1. Thus, it is possible that PU.1 is not the only factor involved in antagonising GATA3, TCF1 and Notch, and further work should define better the roles of other genes in this initial period.

| $p_1$ | $p_2$ | $p_3$ | $p_4$ | $p_5$ | $p_6$ | $p_7$ | $p_8$ | $p_9$ | $p_{10}$ |
|---|---|---|---|---|---|---|---|---|---|
| 0.10 | 1.00 | 5.00 | 1.50 | 0.01 | 0.50 | 0.70 | 0.50 | 1.00 | 0.20 |
| $p_{11}$ | $p_{12}$ | $p_{13}$ | $p_{14}$ | $p_{15}$ | $\gamma_R$ | $\gamma_T$ | $\gamma_G$ | $\gamma_P$ | $\gamma_X$ |
| 2.50 | 2.60 | 2.00 | 1.00 | 0.01 | 0.15 $h^{-1}$ | 0.15 $h^{-1}$ | 0.23 $h^{-1}$ | 0.06 $h^{-1}$ | 0.02 $h^{-1}$ |

TABLE S1 PARAMETER VALUES FOR THE TRANSCRIPTION LEVEL MODEL.

The parameter values for the simplified computational model for *Bcl11b* regulatory system transition from chromatin closed to open state used for simulation results shown in Fig. 8D are shown in Table S2.

| $p_{16}$ | $p_{17}$ | $p_{18}$ | $p_{19}$ | $p_{20}$ |
|---|---|---|---|---|
| 0.05 | 0.05 | 0.01 | 0.20 | 0.10 |

TABLE S2 PARAMETER VALUES FOR THE EPIGENETIC LEVEL MODEL.

**Population Model**

In order to take population dynamics into account in a multi-level model, we developed population models with parameters extracted from the CTV data. As CTV staining intensity provides information about the number of divisions that an individual cell has gone through, it allows us to build generation profiles for each of the assessed cell type groups (DN1 and DN2a) and at different time points of measurements. These profiles exhibit dispersion of the generation distributions in time.

**A simplified model.** To assess if this dispersion can be explained by proliferation, which is uniform for all generations, we first developed a population proliferation model in which the division rate of a cell does not depend on its generation.

A three-parameter model was employed that gave the best fit of the predicted to measured cell numbers with the substantial different average relative error at the points of measurement. The prediction of DN1 cell numbers differed on average from experimental values by 16.9% at day 2 and by 5.3% at day 3 (Table S3). At day 2 in both groups of cells we observed a large proportion (above 50%) of the cell population not dividing between measurements. At day 3 this proportion was lowered to about 20%, with most of the cells (around 50%) having divided twice. This demonstrates that the cells after sorting and seeding into the cell plate culture experienced an initial slowdown in proliferation rate to a value smaller than 1/24h but recovering to the rate of about 1/12h at later times.

The results of this simple population model of cell proliferation suggest that the dispersion of the cell proportions among different generations is not uniform either between different cell types (DN1 or DN2a) or for different measurement time points (day 2, day 3) within the same cell group.

In order to estimate the cell cycle lengths in a population of immature thymocytes as they progress through their development, we first devised a simple population model of cell division fitted to experimental CTV data. This "null model" assumes that a cell can divide between measurement time points from 0 to 3 times. This assumption matches our observations from confocal imagining of the cell cultures. The proportions of the cells in the population undergoing 1 to 3 divisions are denoted *a, b* and *c* respectively. These parameters have to satisfy the relation *a+b+c < 1*. The proportion of cells not dividing between measurements points is given by *1-a-b-c*, which exhausts all the cases considered in the model. We also hypothesize in this "null model" that these proportions are uniform through all the cell generations in the population. Correctness of this working hypothesis will be assessed from the results of the model. This means that number of cells in generation $G_i^{n+1}$ at time $t_{n+1}$ is given in terms of generations at previous time $t_n$ by

$$G_i^{n+1} = (1 - a - b - c)G_i^n + 2aG_{i-1}^n + 4bG_{i-2}^n + 8cG_{i-3}^n$$

Measuring the goodness of the fit to CTV data is given by the relative error (see Table S3).

| | DN1 day 2 | DN1 day 3 | DN2a day3 | DN2a day 3 |
|---|---|---|---|---|
| **Relative error of distribution fit** | 16.9% | 5.3% | 17.4% | 4.2% |

TABLE S3 RELATIVE ERROR FOR THE MODEL ABOVE FITTED TO THE CTV DATA.

This demonstrates that the cells after seeding into the cell plate culture experience initial slowdown in proliferation rate to a value smaller than 1/24h recovering the rate of about 1/12h at later times.

The results and the outputs of this population "null model" of cell proliferation suggest that the dispersion of the cell proportions among different generations is not uniform either between different cell types (DN1 or DN2a) or for different measurement time points (day 2, day 3) within the same cell group.

The CTV data provides complete untruncated distributions of generations 0 to 6 for two groups of cells (DN1 and DN2a cells) for days 2, 3 and 4. This allows us to predict with the model distributions at day 3 given data at day 2 and to predict distributions at day 4 given data at day 3. Comparison of these predictions to the actual data gives us best fit parameters of population level cell proliferation at given time point (Table 2). In this way the dispersion of cell proportions among different generations can be included. These parameters were found by global minimisation of the normalised mean error measure between predicted and measured cell numbers in generations.

| generation | μ | σ |
|---|---|---|
| 0 | 34 | 13 |
| 1 | 15 | 5 |
| 2 | 13 | 5 |
| 3 | 12 | 4 |
| ≤ 4 | 12 | 3 |

TABLE S4 AVERAGE AND STANDARD DEVIATION PARAMETERS FOR CELL CYCLE LENGTH DISTRIBUTIONS FOR EACH GENERATION FITTED TO EXPERIMENTAL MEASUREMENTS (TIMES IN HR).

#### Parameters Epigenetic Level – Multi-scale Model

Since the previously presented "simple" population based model assumed cell cycle lengths independent from cell generations and required different parameter sets for each data time point, we wanted to see if now we can construct unified model of cell proliferation consistent with the data by assuming that the cell cycle length is a function of the cell generation. As such, the parameters of the cell cycle length normal distributions were chosen independently for each cell based on its generation and fitted globally to the CTV data in Table S4. In order to take into account, the effect of cell division on the amounts of epigenetic factors inside a cell, we implemented a Bcl11b regulation region collaborative epigenetic model Fig. S7 with parameters shown in Table S5.

| $k_1$ | $k_2$ | $k_3$ | α | β | γ | δ | ε |
|---|---|---|---|---|---|---|---|
| 0.28 | 0.20 | 0.20 | 0.002 | 0.002 | 0.0005 | 0.0005 | 0.002 |

TABLE S5 PARAMETER VALUES FOR THE COLLABORATIVE EPIGENETIC LEVEL MODEL

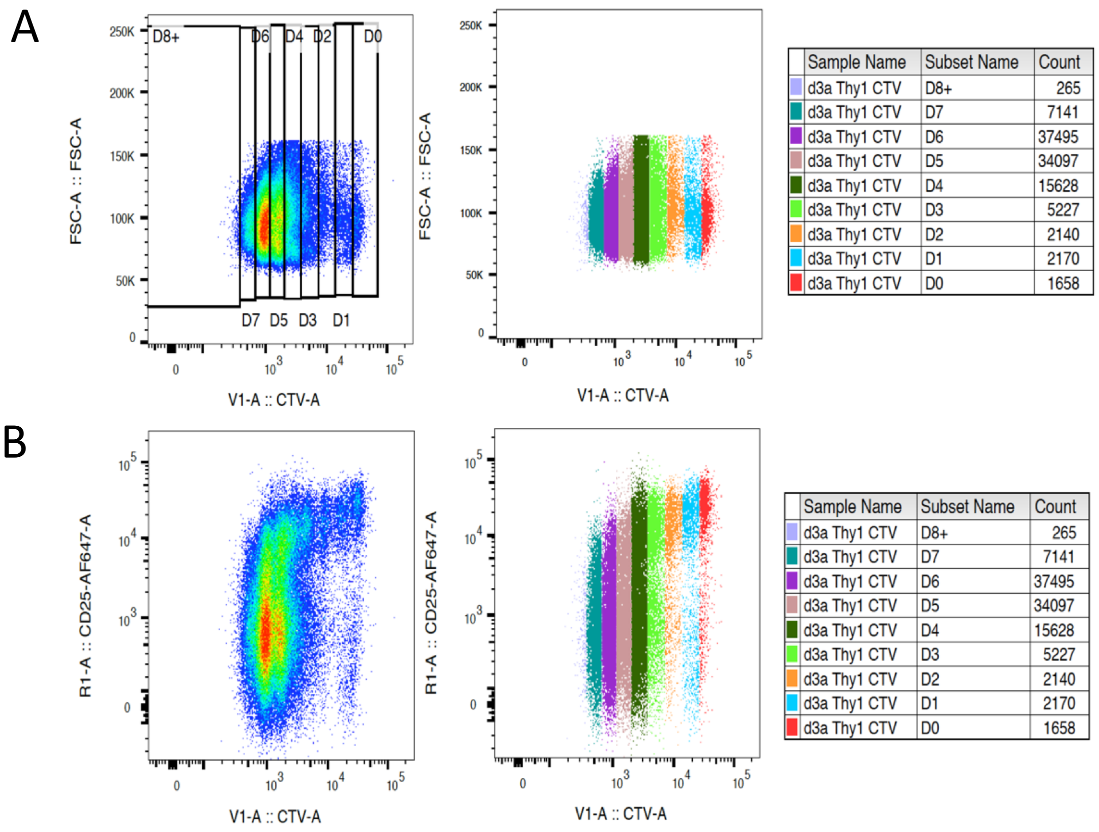


| | Sample Name | Subset Name | Count |
|---|---|---|---|
| | d3a Thy1 CTV | D8+ | 265 |
| | d3a Thy1 CTV | D7 | 7141 |
| | d3a Thy1 CTV | D6 | 37495 |
| | d3a Thy1 CTV | D5 | 34097 |
| | d3a Thy1 CTV | D4 | 15628 |
| | d3a Thy1 CTV | D3 | 5227 |
| | d3a Thy1 CTV | D2 | 2140 |
| | d3a Thy1 CTV | D1 | 2170 |
| | d3a Thy1 CTV | D0 | 1658 |


| | Sample Name | Subset Name | Count |
|---|---|---|---|
| | d3a Thy1 CTV | D8+ | 265 |
| | d3a Thy1 CTV | D7 | 7141 |
| | d3a Thy1 CTV | D6 | 37495 |
| | d3a Thy1 CTV | D5 | 34097 |
| | d3a Thy1 CTV | D4 | 15628 |
| | d3a Thy1 CTV | D3 | 5227 |
| | d3a Thy1 CTV | D2 | 2140 |
| | d3a Thy1 CTV | D1 | 2170 |
| | d3a Thy1 CTV | D0 | 1658 |


**Supplementary Figure S1.**

Cell division determinations using to Cell Trace Violet (CTV) levels in flow cytometric plots of control cells (for Figure 2 CTV data). Unfractionated DN thymocytes were CTV stained and placed in OP9-DL1 culture for 4 days before harvesting, staining with surface differentiation marker antibodies, and FACs analysis (see Methods). CTV levels are diluted by dividing between daughter cells with every cell division. (A) Using Forward Scatter (FSC) vs. CTV plots, gates were drawn as shown, defining how many cell divisions each cell had experienced based on CTV ranges. Cells within each gate were assigned cell division numbers and color coded (right plot and legend): cells that had divided 0 times (D0, red) through 7 times (D7, teal) were distinguished and cells with CTV values below D7 were counted as at least 8 divisions (D8+, lilac). (B) FACs plots of the same cells but showing the relationship between CD25 and CTV in the developing DN cells. For total DN cells (mostly DN3 and DN4), CD25 generally declines/stays off with cell division.

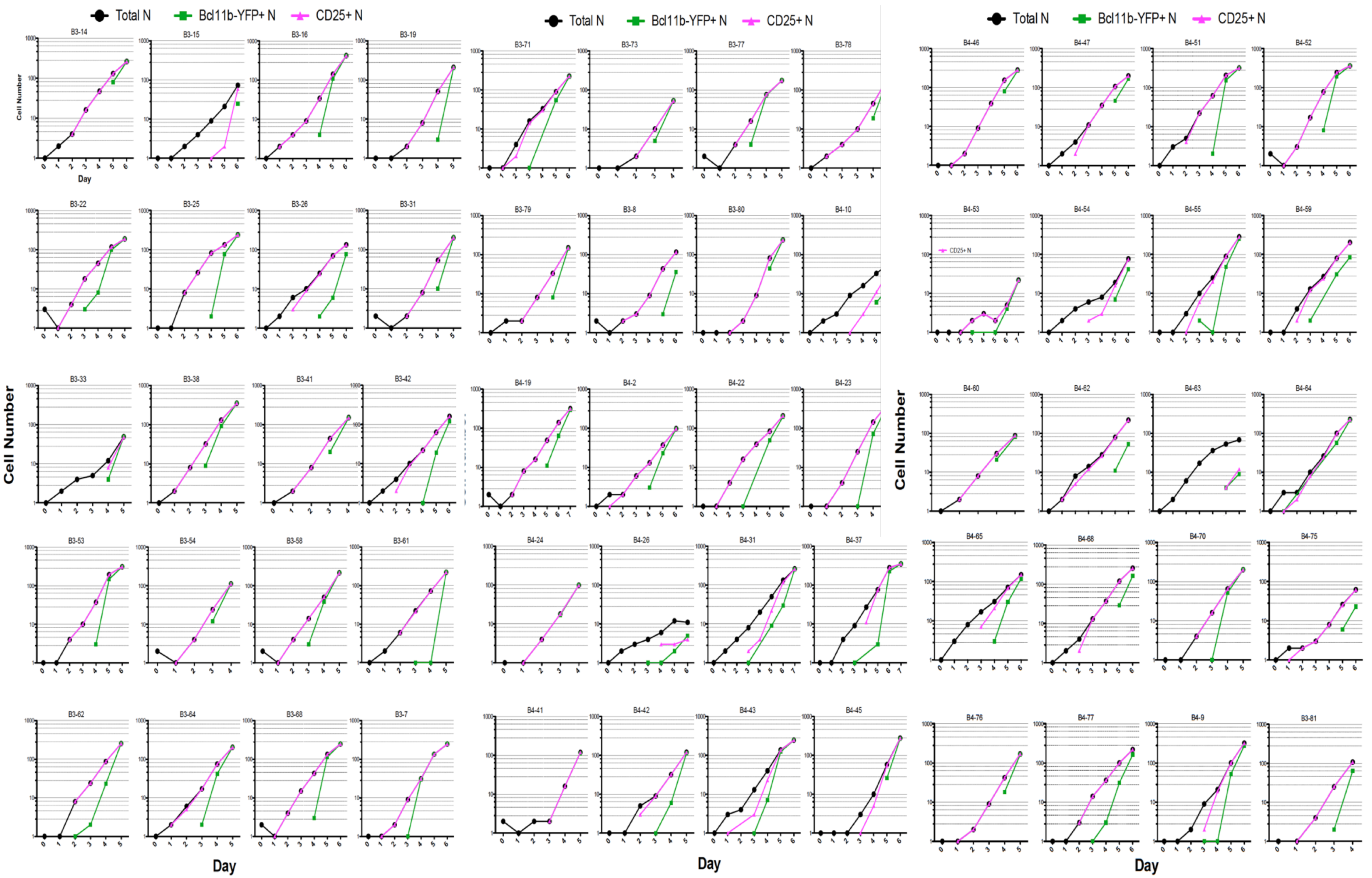


**Supplementary Figure S2.**

Clonal variations in proliferation and differentiation rates. Plots are of cell numbers (log scale) and numbers of CD25 and Bcl11b-YFP positive cells by day for 60 individual clones that clearly entered the T-cell pathway, as determined by turning on CD25 by day 6 (see Methods) black circles=total cell numbers; pink triangles=number of CD25+ cells; green squares=number of Bcl11b-YFP+ cells; 0 values are not plotted.

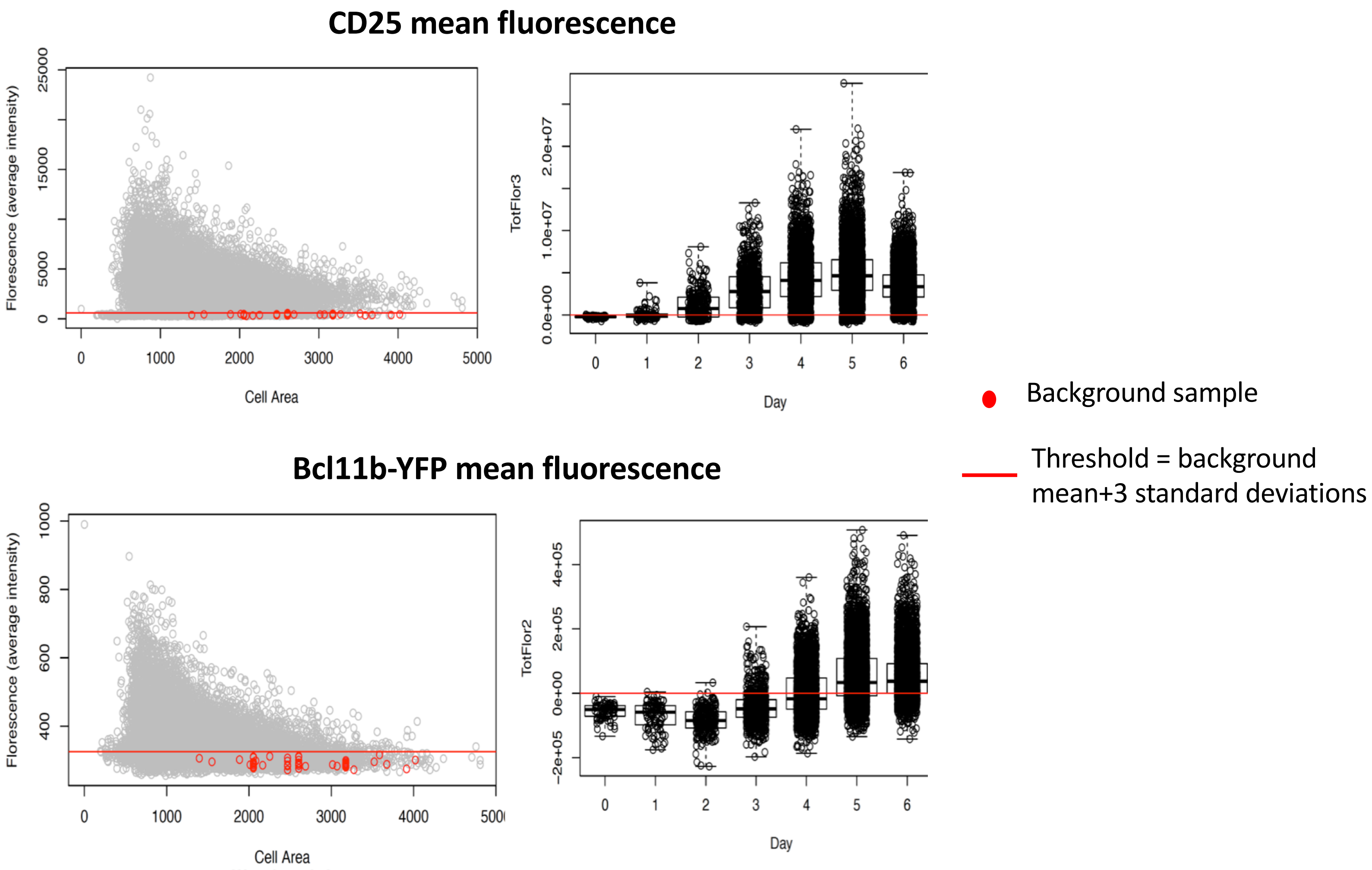


**Supplementary Figure S3**.
Fluorescence threshold level determinations for CD25 and Bcl11b-YFP expression. Plots of fluorescence values for ~13K segmented cells from all ETP clones and all times (left) and fluorescence values for all cells by day (right; median +/- 25th percentile). Background values were taken from 50 samples from 13 wells (red circles) and the average + 3 standard deviations (red line) was selected as the threshold level of expression for scoring a cell as being positive for CD25 or Bcl11b-YFP.

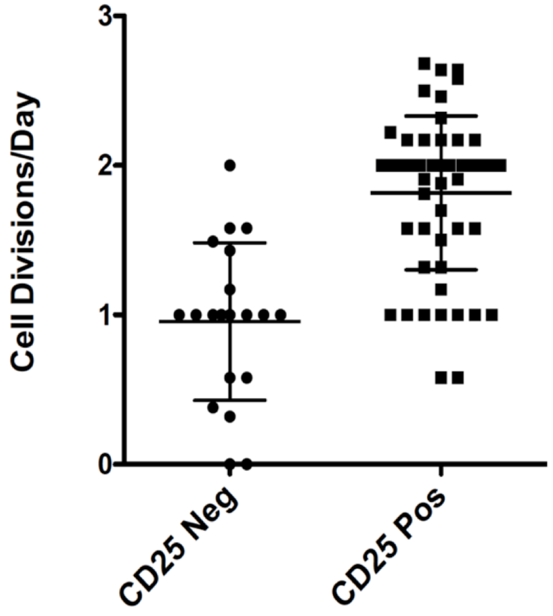


**Supplementary Figure S4.**
The estimated number of divisions/day for clones in which all daughter cells were scored as either CD25 negative (left) or positive (right) for two consecutive days in early cultures, from days 1-3. Cells from clones that had turned on CD25 (DN2) proliferated more rapidly than those that had not turned on CD25 (still DN1) (DN1, n=20; DN2, n=55; p<0.0001 paired t-test).

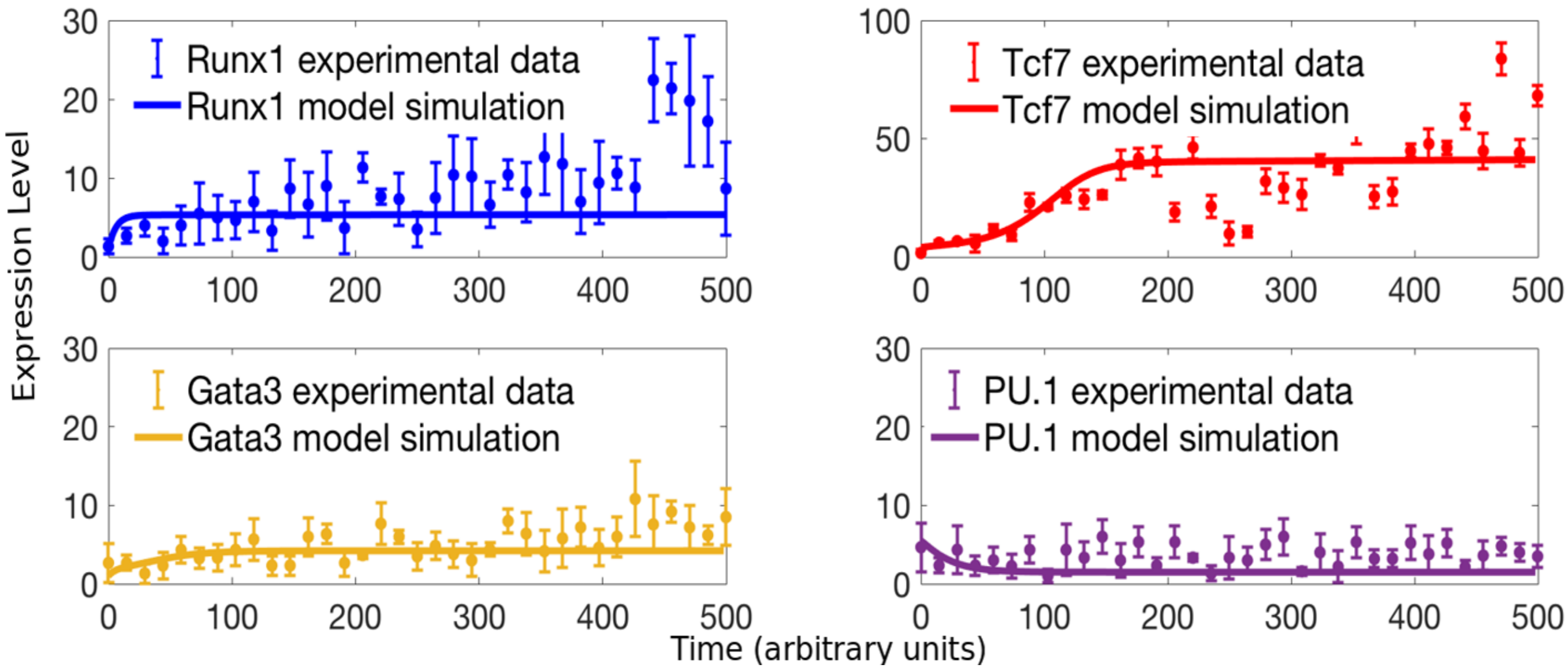


**Supplementary Figure S5.**
Best model fits to pseudo-time-series data. Simulation dynamics from the gene regulatory network model with optimised parameters versus the pseudo-time-series data. The continuous lines depict the model deterministic simulation results while the dots show the mean mRNA count in each cluster obtained from FISH data and the bars show standard deviations.

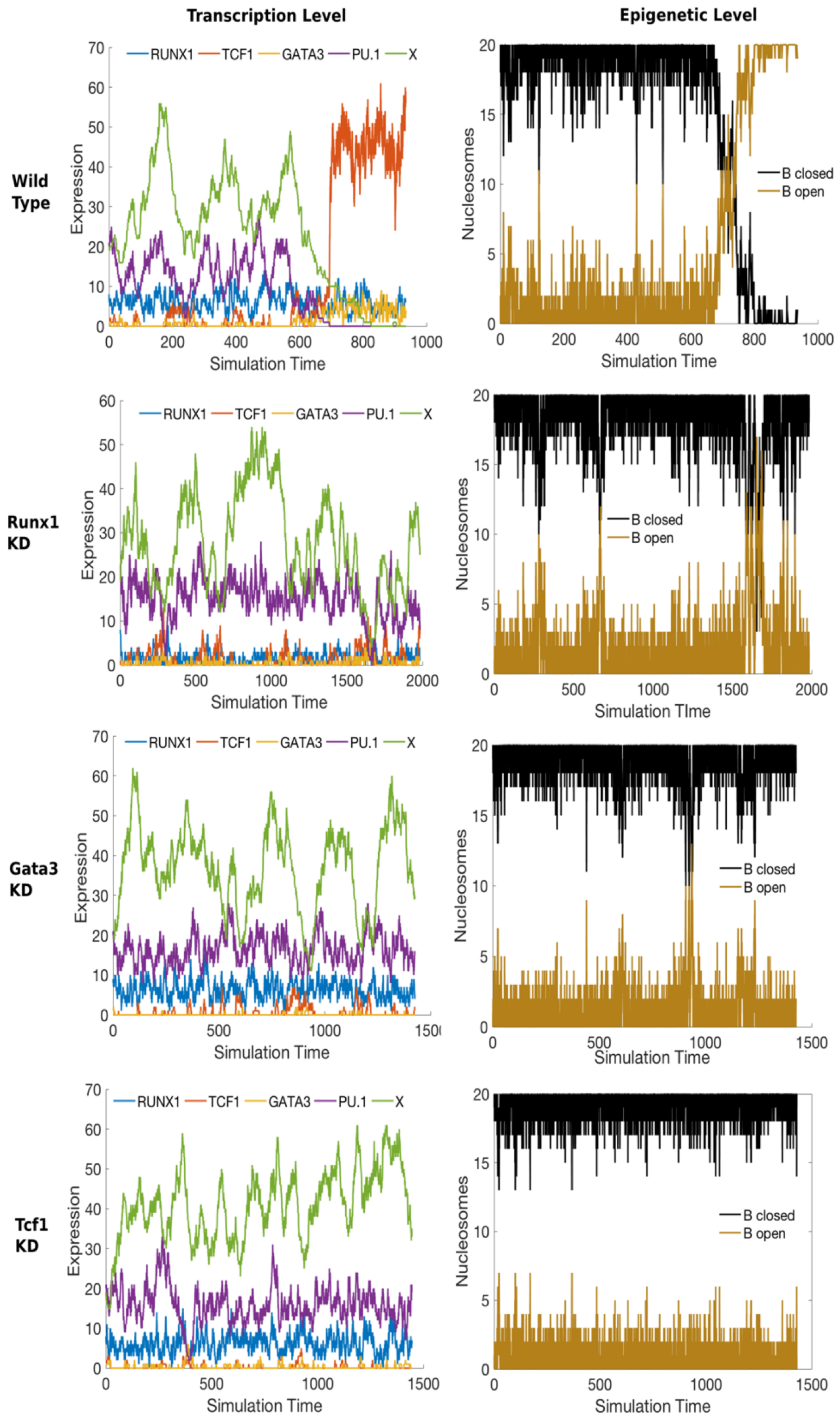


**Supplementary Figure S6.**
Knock-down simulations for DN1 cells. First row shows the wild-type where no gene was knocked-down (K-D). The switch is thrown at the transcriptional level and Bcl11b becomes open. Second, third and fourth rows show Runx1, Gata3 and Tcf1 K-D simulations. The T-cell factors do not get expressed and Bcl11b remains closed.

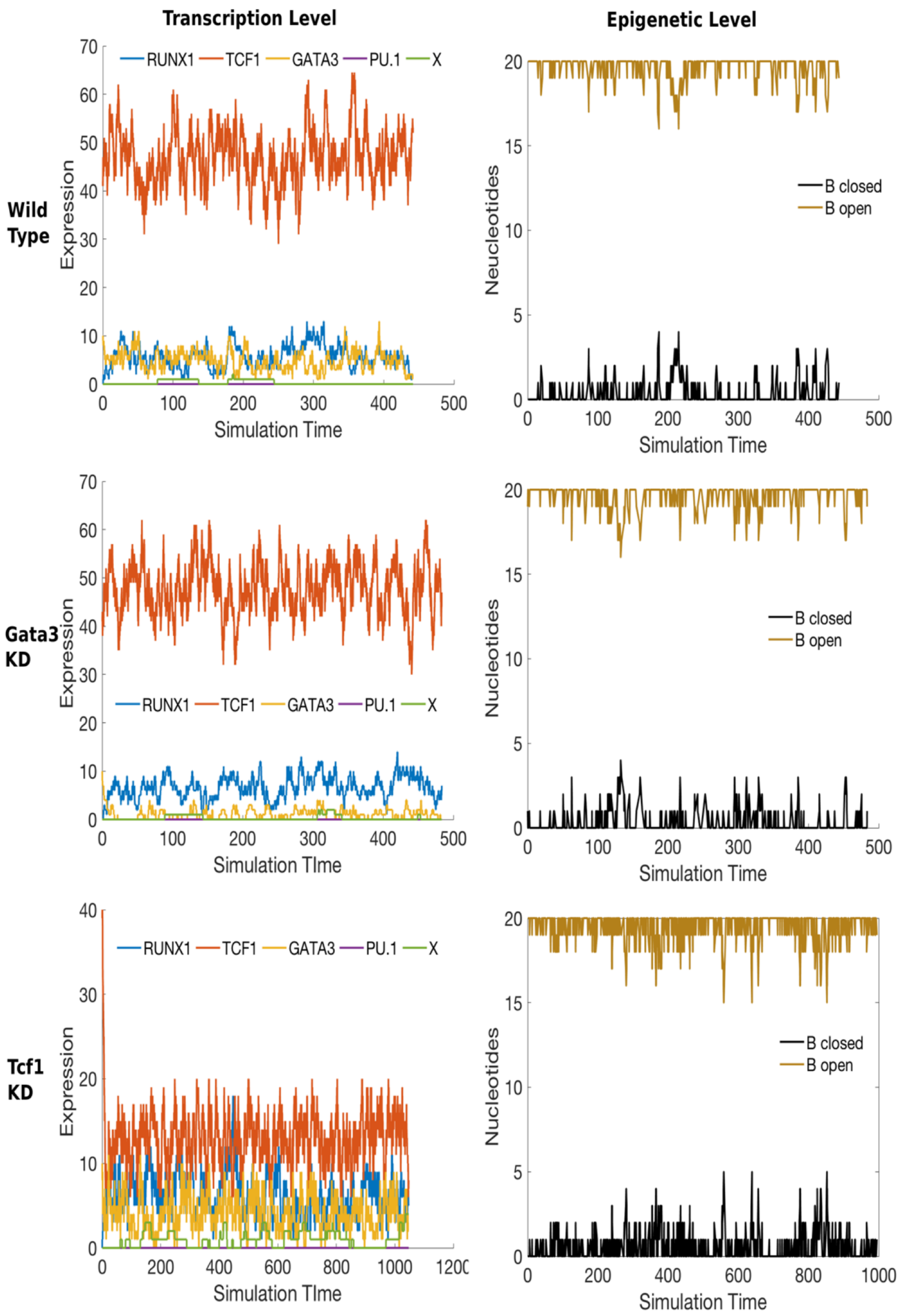


**Supplementary Figure S7.** Knock-down simulations for DN2 cells. First row shows the wild-type, where no gene was knocked-down. The T-cell factors are high and X is low, thus Bcl11b is open. Second and third rows show Gata3 and Tcf1 K-D simulations. Bcl11b remains open, even though Gata3 and Tcf1 were K-D.

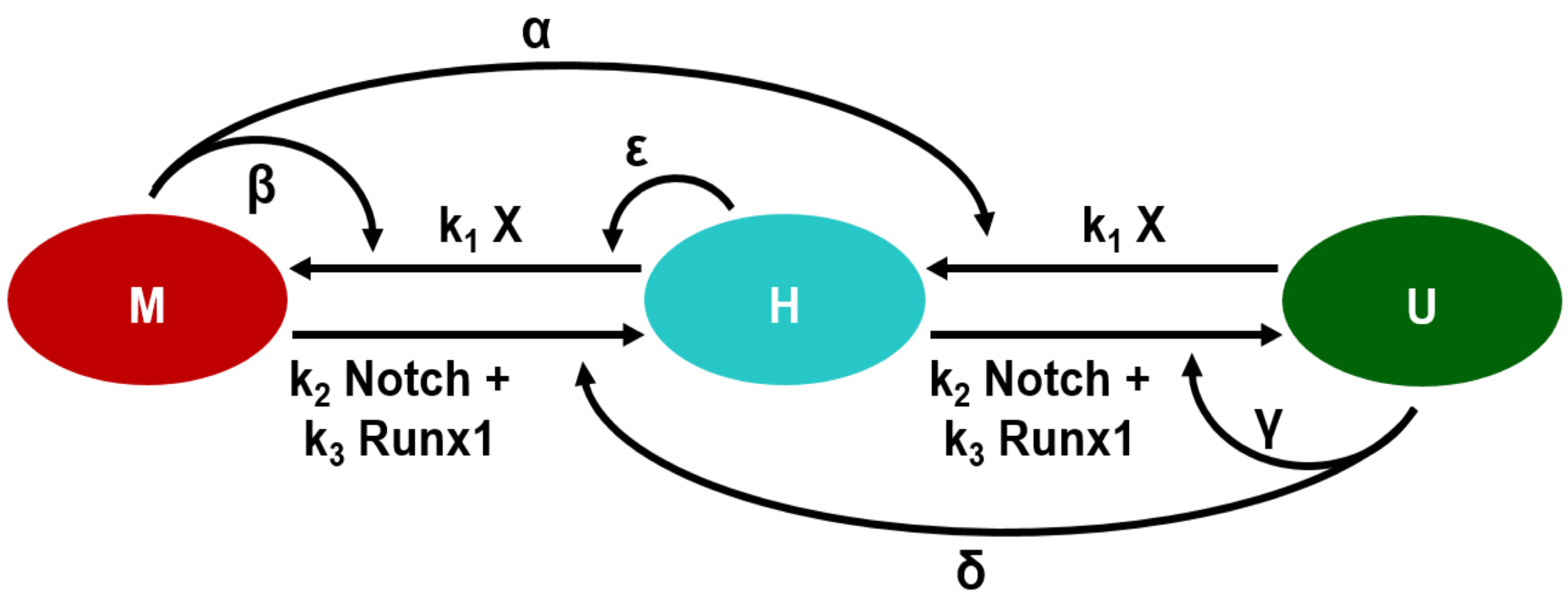


**Supplementary Figure S8.**
Collaborative model used to simulate the demethylation of the Bcl11b regulatory region. CpG sites can be methylated (M) -- closed state, hemi-methylated (H) -- intermediate state or unmethylated (U) -- open state.
